# Structural disruption of exonic stem-loops immediately upstream of the intron regulates mammalian splicing

**DOI:** 10.1101/292003

**Authors:** Kaushik Saha, Whitney England, Mike Minh Fernandez, Tapan Biswas, Robert C. Spitale, Gourisankar Ghosh

## Abstract

Recognition of highly degenerate mammalian splice sites by the core spliceosomal machinery is regulated by several protein factors that predominantly bind exonic splicing motifs. These are postulated to be single-stranded in order to be functional, yet knowledge of secondary structural features that regulate the exposure of exonic splicing motifs across the transcriptome is not currently available. Using transcriptome-wide RNA structural information we show that retained introns in mouse are commonly flanked by a short (≲70 nucleotide), highly base-paired segment upstream and a predominantly single-stranded exonic segment downstream. Splicing assays with select pre-mRNA substrates demonstrate that loops immediately upstream of the introns contain pre-mRNA-specific splicing enhancers, the substitution or hybridization of which impedes splicing. Additionally, the exonic segments flanking the retained introns appeared to be more enriched in a previously identified set of hexameric exonic splicing enhancer (ESE) sequences compared to their spliced counterparts, suggesting that base-pairing in the exonic segments upstream of retained introns could be a means for occlusion of ESEs. The upstream exonic loops of the test substrate promoted recruitment of splicing factors and consequent pre-mRNA structural remodeling, leading up to assembly of the early spliceosome. These results suggest that disruption of exonic stem-loop structures immediately upstream (but not downstream) of the introns regulate alternative splicing events, likely through modulating accessibility of splicing factors.

## INTRODUCTION

To remove introns, mammalian splicing machinery recognizes 5′ (donor) and 3′ (acceptor) splice sites (SS) flanking the intron, as well as the intervening branchpoint site (BS) and the polypyrimidine tract (PPT) near the 3′-SS, yet the precise mechanisms of recognition remain unclear (1). The functionality of these ‘splice signals’ is dependent not only on their sequences but also on the context in which they are present within the pre-mRNA (2). Exonic segments flanking an intron contain a variety of splicing regulatory elements (3) including exonic splicing enhancers (ESEs) and exonic splicing silencers (ESSs) (4), both of which contain loosely conserved sequences for engagement of different RNA binding proteins and are important for regulated assembly of the spliceosome (5). One of the most studied regulators of ESE-dependent splicing activation is the serine-arginine-rich (SR) family of RNA-binding proteins that contain an N-terminal RNA-binding domain (RBD) and an Arg-Ser-rich (RS) domain at their C-terminus (6). SR-proteins promote constitutive splicing and regulate alternative splicing *in vivo* by binding to ESEs present within a short distance (∼50 nucleotides) upstream and downstream of 5′- and 3′-SS, respectively (7-11). Other *trans*-acting splicing factors have also been implicated in ESE-dependent splicing activation (12).

Recent studies indicate that RNA transcripts can adopt multiple secondary and tertiary structures *in vivo* (13-17), and that these structures modulate binding to partner protein complexes and regulate pre-mRNA processing (18-20). Co-transcriptional RNA folding is also known to be regulated by the rate of transcription elongation in eukaryotes (21), suggesting a coordinated regulation of splicing and transcription elongation (22-26). Although it is postulated that functional ESE and ESS sequences are present on single-stranded RNA (27), little information on the correlation of secondary structure of pre-mRNAs with transcriptome-wide distribution of ESE/ESS sequences for splicing regulation is available.

In this study, we investigate the secondary structure of intron proximal exonic regions. We observe that exonic segments immediately upstream of the 5′-SS display increased base-pairing in retained introns relative to spliced introns of the mouse embryonic stem cell transcriptome. A detailed investigation of these exonic segments to correlate secondary structure, the presence of enhancer/silencer elements, and SR protein binding indicates that the degree of base-pairing within exonic segments immediately upstream of the 5′-SS regulates assembly of the early spliceosome.

## MATERIALS AND METHODS

### Cloning and protein expression

cDNAs of SR proteins were purchased from Open Biosystems and cloned in a T7 promoter-based *E. coli* expression vector (pET24d; EMD Millipore). Proteins contained a hexa-histidine (His_6_) - tag that is removable by *TEV* protease. Proteins were expressed in *E. coli* BL21 (DE3) cells with isopropyl β-D-1-thiogalactopyranoside induction and purified by Ni^2+^-nitrilotriacetate (Ni-NTA) affinity chromatography. The His_6_-tags were removed from all proteins used in our experiments by treatment with His_6_-*TEV* protease overnight at room temperature, and residual uncleaved proteins and His_6_-tagged *TEV* protease were removed by subsequent passage through Ni-NTA resin. Tag-removed proteins were further purified by size-exclusion chromatography (Superdex 75; GE Healthcare Lifesciences). Two functional variants of SR proteins used in our experiments were the RNA binding domain (RBD) of SRSF1 (amino acids 1-203)(28) and RBD of SRSF2 (amino acids 1-127) in chimera with fully phosphomimetic RS domain of SRSF1 (amino acids 197-246) in which all serine residues were replaced with glutamate (29). All purified proteins were confirmed to be RNase-free by incubating a small aliquot of the purified protein with a long RNA (e.g. *β-globin* RNA) overnight at room temperature and analyzing the RNA quality by urea-PAGE after phenol extraction.

### Pre-mRNA constructs

The mouse pre-mRNA constructs with retained introns used in this study are *2610507B11Rik* IVS17 (192+127+203) (chr11:78272778-78273299, mm10) and *Amt* IVS1 (104+102+427) (chr9:108296922-108297295, mm10). The model pre-mRNA substrates used are human *β-globin* IVS1 (109+130+204) and Adenovirus 2 major later transcript IVS1 (*AdML*) (65+123+50). The lengths of exon-intron-exon segments of pre-mRNA constructs are provided within parenthesis.

### Electrophoretic mobility shift assay (EMSA)

Uniformly radiolabeled pre-mRNA was synthesized with [*α-P*^*32*^]*UTP* (3000 Ci/mmol; 10 µCi/µl) by *in vitro* (run off) transcription driven by T7 RNA polymerase (New England Biolabs), Next, the DNA template was digested with 2 units of DNase I (New England Biolabs) for 1 h at 37 °C and the RNA was desalted twice by Illustra Microspin G-25 columns (GE Healthcare Life Sciences). Method for EMSA was modified from a previously published protocol (30). ∼10 pM radiolabeled pre-mRNA was incubated with an SR protein for 20 min at 30 °C in 20 mM HEPES-NaOH (pH 7.5), 250 mM NaCl, 1 mM DTT, 2 mM MgCl_2_, 1 M urea, 20% glycerol, and 0.3 % polyvinyl alcohol (P-8136, Sigma) in 15 µl volume. Reaction products were resolved on 4 % (89:1) polyacrylamide gels containing 2.5 % glycerol and 50 mM Tris-glycine buffer. Gels were run at 250 V for 90 min at 4 °C, dried, and analyzed by phosphorimaging.

### <u>S</u>elective 2′-<u>h</u>ydroxyl <u>a</u>cylation analyzed by <u>p</u>rimer <u>e</u>xtension (SHAPE)

The SHAPE assay was performed broadly following a protocol published before (31). Denatured and renatured *β-globin* pre-mRNA (25 nM) was incubated with SRSF1 (750 nM), SRSF2 (250 nM), or an equal volume SR protein storage buffer in the abovementioned EMSA condition (without glycerol) for 15 min. Similarly, denatured and renatured *AdML* pre-mRNA (25 nM) was incubated with SRSF1 (250 nM). Freshly prepared N-Methylisatoic anhydride (NMIA, Aldrich) in dimethyl sulfoxide (DMSO, Sigma) was added to the test reactions at 2 mM (final concentration), and the reaction was allowed to proceed for 15 min at 30 °C and then 15 min at room temperature. Equal volume of DMSO was added to the control reactions. Reaction mixtures were treated with proteinase K at 37 °C for 10 min (New England Biolabs) and the RNA samples were purified by RNeasy Mini Kit (Qiagen). Reverse transcription reaction was carried out using MuLV Super-RT reverse transcriptase (Biobharati Life Science, India) using manufacturer’s protocol at 50 °C. cDNAs were precipitated with ethanol, resuspended in deionized formamide containing bromophenol blue, xylene cyanol, and 0.5 mM EDTA, and separated in 12 % and 8 % 7 M urea sequencing gels of 0.35 mm thickness. In 8 % gels, the same samples were loaded a second time into empty lanes when xylene cyanol of the first load migrated 12.5 cm. Electrophoresis was continued till xylene cyanol from the second load migrated 28 cm. We could resolve up to 125 nucleotides with each primer by this method. The individual bands were identified and quantified using Fiji (32). Where the lanes did not run straight, they were divided into several segments with overlapping regions, with each individual segment being straight. The quantified band intensities within the overlapping regions were used to normalize the values between consecutive segments. The independent values of a primer series from different gel runs (i.e. 12 % and 8 %) were also normalized. The band intensity values of the DMSO-treated control RNA were subtracted from those of NMIA-treated pre-mRNAs to obtain the raw SHAPE reactivity profiles. The raw reactivity data were processed as described in the earlier work (31). Briefly, all raw SHAPE reactivity values for a primer series were divided by the average of the top 10 % values excluding the high-value outliers to obtain the processed values. Raw SHAPE reactivity values that were ≥ 1.5 times the median of all values for a primer series – capped at top 5 % of all band intensity values in each primer series as the number of bands obtained with each primer was small, about 100-150 – were considered outliers. All negative values were considered to be zero. Maximum values were capped at 3. For *β-globin*, we also processed the raw SHAPE-reactivity by 90 % Winsorization for comparison to the trimming-based method described above. The processed values were analyzed with the ‘RNAstructure’ (33) software, which considered these values as pseudo-free energy of folding to generate the secondary structure models. Two variable parameters of RNA folding, the slope and the intercept, were set to 2.6 and -0.8, respectively. The RNA secondary structure models were drawn using VARNA (34). ΔSHAPE-reactivity for SR protein-bound RNA was calculated by subtracting the SHAPE reactivity of each nucleotide of the protein-free RNA from that of the corresponding nucleotide of the SR protein-bound RNA.

The primers used for reverse transcription of *β-globin* were 5′ACGTGCAGCTTGTCACAGTG (*βgRT1*), 5′TTTCTTGCCATGAGCCTTC (*βgRT2*), 5′AGTGGACAGATCCCCAAAG (*βgRT3*), 5′GGAAAATAGACCAATAGGC (*βgRT4*), 5′TTCTCTGTCTCCACATGCC (*βgRT5*), and 5′AACTTCATCCACGTTCACC (*βgRT6*). For reverse transcription of the EH3+4 mutant of *β-globin, βgRT6* was replaced with *EH-RT6* (5′CCAACTTCATGCTGGTGACG). Reverse transcription of *AdML* was performed with primers: 5′AAGAGTACTGGAAAGACCGC (*AdRT1*), 5′GGACAGGGTCAGCACGAAAG (*AdRT2*), and 5′CAGCGATGCGGAAGAGAGTG (*AdRT3*).

### *In vivo* splicing assay

*In vivo* splicing assays with minigene constructs were performed in *HeLa* cells grown in a six-well plate (9 cm^2^ surface area) transfected with 1 µg of the purified plasmid. RNA was harvested 36 h post transfection using Trizol reagents (Thermoscientific) following manufacturer’s instructions, treated with RNase-free DNase I (New England Biolabs), and purified by phenol:chloroform extraction and ethanol precipitation. cDNA was synthesized in a 20 µl volume using 100 ng RNA with MuLV Super RT reverse transcriptase (Biobharati Life Science, India) at 50 °C following manufacturer’s instructions. 1 µl of this cDNA was used in a 25 µl reaction volume for PCR. Non-saturating PCR products were analyzed in 2 % agarose gels. Densitometric estimation was carried out by Fiji (32). The splicing efficiency (% spliced) was calculated using the formula: (mRNA/(mRNA + pre-mRNA + aberrant product))/(WT mRNA/(WT mRNA + WT pre-mRNA))*100. Each splicing assay was repeated three times. All substrates were expressed *in vivo* under the control of a CMV promoter (pcDNA3.1) since it represents a strong promoter (transcription rate ∼6-7 mRNAs per hour) near the median strength of endogenous mammalian promoters (35,36). New England Biolabs (Ipswich, MA) DNA ladder 100 bp is used throughout the study and all size markers are annotated in order to indicate the size of the RT PCR products.

### *In vitro* spliceosome assembly assay

Nuclear extract was prepared from *HeLa* cells as published (37). Spliceosomal complex assembly assay was carried out as described before (38).

### RNA-seq data analysis

icSHAPE data was generated from publicly available raw data (15). The datasets polyA(+) icSHAPE DMSO biological replicates (Gene Expression Omnibus accession # GSM1464037, GSM1464038), polyA(+) icSHAPE *in vitro* NAI-N_3_ biological replicates (GSM1464039, GSM1464040), and polyA(+) icSHAPE *in vivo* NAI-N_3_ biological replicates (GSM1464041, GSM1464042) were used. The icSHAPE enrichment score was calculated as in (15) using previously published icSHAPE pipeline (https://github.com/qczhang/icSHAPE). Briefly, the difference between the reverse transcription stops in the treated sample and the control sample was divided by base density in the control sample. Outliers were removed by 90 % Winsorization and the scores were then scaled from 0 to 1. This method tacitly integrates the biological variation between replicates into the icSHAPE enrichment score but does not generate separate icSHAPE enrichment score for individual replicates. Since NAI-N_3_ adducts formed at flexible 2’-OH group of the RNA is quickly hydrolyzed, only reasonably stable RNA structural features are likely captured by NAI-N_3_.

All exon-exon and exon-intron boundaries were identified using annotations in the GENCODE release M12. Only transcripts annotated to contain retained introns were searched for exon-retained intron boundaries. To identify the position of the retained intron, exon start and stop positions from retained intron transcripts were compared to those of coding transcripts originating from the same gene. Exons from retained intron transcripts that spanned two sequential exons in a related coding transcript were considered to contain introns, and the exon-retained intron boundaries were defined as the 3′ end of the upstream exon and the 5′ end of the downstream exon in the coding sequence. A 70-nt long region upstream of the 5′ boundary and downstream of the 3′ boundary was extracted for analysis. For comparison, 70-nt upstream and downstream of 10,000 randomly selected annotated exon-exon junctions in coding transcripts were analyzed. For both sets, transcripts with another exon-exon or exon-intron boundary, or a transcript end, within the 70 bp were discarded. The same method was used to isolate the exons downstream of the introns. Custom scripts sjshapePC.py and sjshapePC_3prime.py were used to collect icSHAPE enrichment score upstream and downstream of exon-exon junctions, respectively. intronHunterPC.py and intronHunterPC_3prime.py were used to collect the same for the exonic segments upstream and downstream of retained introns. The scripts are made available at https://github.com/englandwe/NAR_splicing. For standard error of the mean (SEM), we calculated the mean of all enrichment scores at the given position relative to the splice junction of a set of transcripts, with or without retained introns.

### Counting ESE and ESS sequences

Hexamers assigned as ESE, ESS, or neutral sequences in (4) within target (70-nt) and non-target (80-nt) regions were counted using Jellyfish (39). ‘Splice score’ was calculated by summing up the individual scores of ESE and ESS as described before (4).

### *In vivo* DMS-seq

*HeLa* cells transfected with the plasmids were treated with dimethyl sulfate as described before (40). For isolation of nuclei (41), transfected and DMS-treated cells were washed with ice-cold phosphate-buffered saline (PBS), scraped cells were centrifuged for 30 s at 1000 g, the supernatant was removed, cells were resuspended in 10 mM HEPES 7.9, 1.5 mM MgCl_2_, 10 mM KCl, 0.5 mM DTT, and 0.05 % Nonidet P-40 and kept on ice for 10 min with occasional flicking. Then the suspension was centrifuged for 10 min at 730 g at 4 °C. The pellet was resuspended in Trizol (Thermoscientific) and RNA was isolated following manufacturer’s protocol. Reverse transcription of *β-globin* constructs was performed with *βgRT5* (nested in the intron ensuring production of the cDNA synthesis from the pre-mRNA and not the mRNA). Reverse transcription and densitometric analyses were carried out as in SHAPE. For densitometric measurements, background of the images was subtracted.

### Statistical analyses

The statistical significance of splicing assay results was calculated using one-sample t-test. *p*-values for icSHAPE and ESE/ESS analyses are from Welch’s two-sample t-tests, with Bonferroni correction; the R-scripts are uploaded onto github (https://github.com/englandwe/NAR_splicing).

### Calculation of ‘probability of being unpaired’ (PU) values by MEMERIS, RNAplfold, and LocalFold

‘Probability of being unpaired’ or PU values for each hexamer within the given sequences were calculated using MEMERIS (27), RNAplfold (42), as well as LocalFold (43). MEMERIS uses RNAfold algorithm (44) to fold the input sequence. For each motif, we iterated MEMERIS calculations 20 times by varying the maximum lengths of the regions flanking the hexamer of interest from 11 to 30 nt (input sequence = 11-30 + hexamer + 11-30) and finally generated an average PU value for each motif. Thus, MEMERIS uses RNAfold to determine local RNA structure. We have made two corrections in the ‘Getsecondarystructurevalues.pl’ script of MEMERIS: We changed the regular expression to allow for multiple spaces preceding the free energy value in the RNAfold output to accommodate current RNAfold output formatting (lines 145 and 200), and modified output text formatting to handle non-numerical return values if RNAfold output is not properly recognized (at line 377). We have uploaded the revised script on github (https://github.com/englandwe/NAR_splicing).

On the other hand, RNAplfold (42) also determines local structure in sliding windows (W) by imposing a maximum base-pair span (L; L≤ W) and calculates the probability of occurrence of a specific base-pair over all folding windows containing the sequence interval with the base-pair. RNAplfold was used with two sets of parameter for calculating PU values of individual hexamers: with L=W=70 and with L=70, W=120 (W=L might cause unusually high base pairing probabilities of long-range base pairs, which could be resolved by setting W=L+50 (43)).

LocalFold uses RNAplfold algorithm to fold the RNA but corrects a prediction bias at each artificial border of the sliding window introduced by RNAplfold by ignoring the outlier values of 10-nt at the window borders but including them in the structure. We used L = 70 and W = 120 for LocalFold calculations.

For transcriptome-wide analysis, we kept up to a 30-nt sequence flanking the 70-nt ‘target region’ (see Results). The flanking 30-nt region from the transcript was collected using custom scripts flankFetch_2way*.py, which are available at https://github.com/englandwe/NAR_splicing.

## RESULTS

### Exonic segments immediately upstream of retained introns are highly base-paired

To assess structural flexibility of exons flanking mammalian splice sites, we analyzed structural profiles of mRNA using ‘*in vivo* click selective 2′-hydroxyl acylation and profiling experiment’ (icSHAPE) data for the mouse embryonic stem cell transcriptome (15). icSHAPE identifies single-stranded nucleotides by measuring 2’-hydroxyl flexibility. mRNAs used in the study are poly-A purified and thus have undergone processing. A schematic of the *in vivo* (reaction performed in cell) and *in vitro* (reaction performed with purified RNA) icSHAPE experiments is shown in Figure 1A. *In vivo* the ribonucleotide flexibility is governed by both base-pairing and by interactions with proteins; *in vitro* only the degree of base-pairing influences the readout. Both *in vitro* and *in vivo* icSHAPE data indicate low average reactivities of nucleotides in exonic regions immediately upstream of the 5′-SS preceding retained introns, suggesting that these segments exhibit a higher degree of base-pairing (Figure 1B). In contrast, equivalent exonic segments that precede spliced-out introns display a higher average icSHAPE reactivity. A similar study of the exonic regions (20-50 nt long) immediately downstream of the retained introns reveal higher icSHAPE reactivities compared to similar regions downstream of spliced introns (Figure 1C).

**Figure 1.**
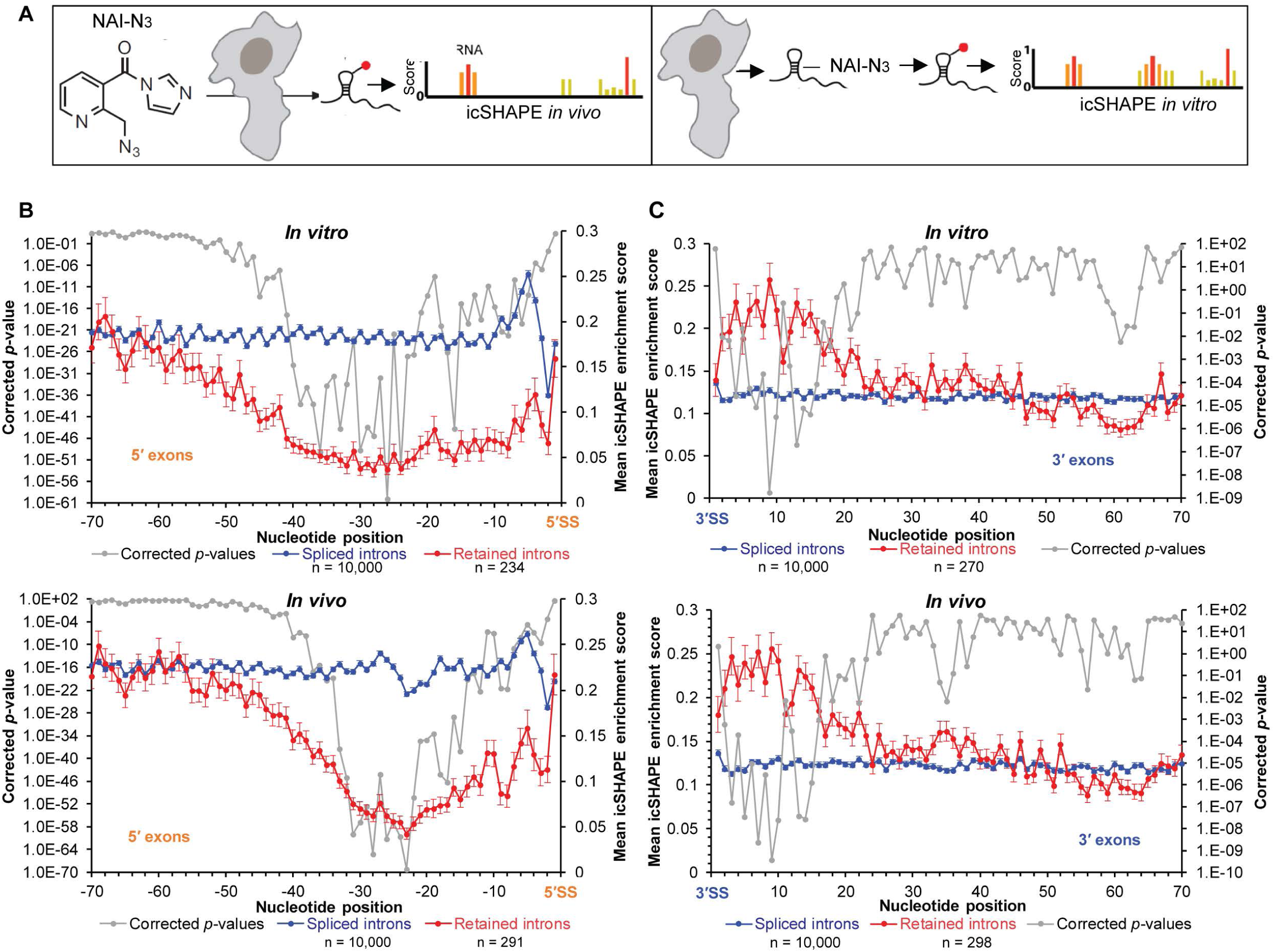
icSHAPE analyses of mRNA from mouse embryonic stem cells. (A) Schematic of *in vivo* and *in vitro* icSHAPE analyses. (B) *In vitro* (top panel) and *in vivo* (bottom panel) icSHAPE enrichment score for 70-nt region upstream of 5′-SS in 10,000 exon-exon junctions with spliced introns (blue line) and 234/291 (*in vitro/in vivo* dataset with 219 shared) exon-intron junctions with retained introns in polyadenylated mRNAs (red line); error bars indicate standard error of the mean indicating variation in icSHAPE enrichment score at each position among the transcripts in each group (retained or spliced); *p*-values (corrected) for icSHAPE reactivity at individual nucleotide positions are indicated with a grey line. (C) *In vitro* (top panel) and *in vivo* (bottom panel) icSHAPE data for 70-nt region downstream of 3′-SS in 10,000 exon-exon junctions with spliced introns (blue line) and 270/298 (*in vitro/in vivo* dataset with 257 shared) intron-exon junctions with retained introns in polyadenylated mRNAs (red line), corrected *p*-values (grey line).

We estimated the G/C content of exonic segments and observed an enrichment in exons upstream of retained introns compared to those upstream of spliced introns (Supplementary Figure S1A). However, no statistically significant difference was observed in the GC content of exonic segments downstream of the retained introns compared to the spliced introns (Supplementary Figure S1B).

Then we calculated PU (probability of being unpaired) values of the exonic segments flanking the retained and spliced introns (described in the Method section). Plots showing the mean PU values per hexamer obtained by MEMERIS, RNAplfold (L=W=70), RNAplfold (L=70, W=120), and LocalFold (L=70, W=120) of the spliced (Supplementary Figure S1C, D) and retained (Supplementary Figure S1E, F) introns across the transcriptome (*in vivo* dataset) are provided. The mean PU value per nucleotide was obtained by averaging the PU values of each hexamer overlapping an individual nucleotide. The six nucleotides of each hexamer were identified based on the fact that the RNAplfold output provides hexamer end-positions and MEMERIS output hexamer start positions. Plots representing the mean PU values per nucleotide are provided in Supplementary Figure S1G, H, I, J. The PU values obtained with RNAplfold (L=70, W=120) and LocalFold (L=70, W=120) appeared to be almost identical. A moderate correlation (r ≈ 0.7) was observed between mean icSHAPE enrichment score and mean PU values of the 70-nt long ‘target region’ upstream of the retained introns obtained by RNAplfold and LocalFold (Supplementary Table S1). The PU values of the other segments obtained with any method did not exhibit moderate or better (r ≥ 0.7) correlation with the mean icSHAPE enrichment score (Supplementary Table S1).

We next investigated variations in size, distribution, and types of the retained introns. Intron size ranges between 67 and 15,574 nucleotides and distribution also varies widely. We looked for the presence of ‘exitrons’ (a small and unique class of retained introns present within protein-coding exons (45), the splicing of which inhibits functional protein synthesis), and observed four human exitron homologues in the *in vitro* mouse retained intron dataset. We asked if a subset of the retained introns could be ‘detained introns’ (i.e., retained introns with longer half-lives undergoing post-transcriptional splicing (46)), and found that about one-fifth of the retained introns are also reported to be detained introns.

### Exonic loops immediately upstream of the 5′-SS are important for splicing

Since the retained introns are preceded by hybridized exonic segments, we hypothesized that loops in this region might be important for splicing activation. We tested this possibility by studying the structures and splicing activities of two well-studied, constitutively spliced substrates, human *β-globin* IVS1 (47) and Adenovirus 2 major late transcription unit IVS1 (*AdML*) (48). First, we determined secondary structural models of these substrates using *in vitro* SHAPE reactivity, which revealed that both substrates have four loops (loops 1 - 4) in the exonic segment immediately upstream of the 5′-SS (see Supplementary Figure S2A and S2B for SHAPE-derived secondary structure models). We then generated mutants with select loops eliminated (by replacing nucleotides on one strand, allowing it to hybridize to the opposite strand) and named them exon hybridization (EH) mutants. We also generated intron hybridization (IH) mutants with loops downstream of the 5′-SS (within introns) hybridized (Figures 2A, 2D, Supplementary Figure S2A, S2B). The 9-nt of 5′-SS and its three flanking nucleotides were unaltered.

**Figure 2.**
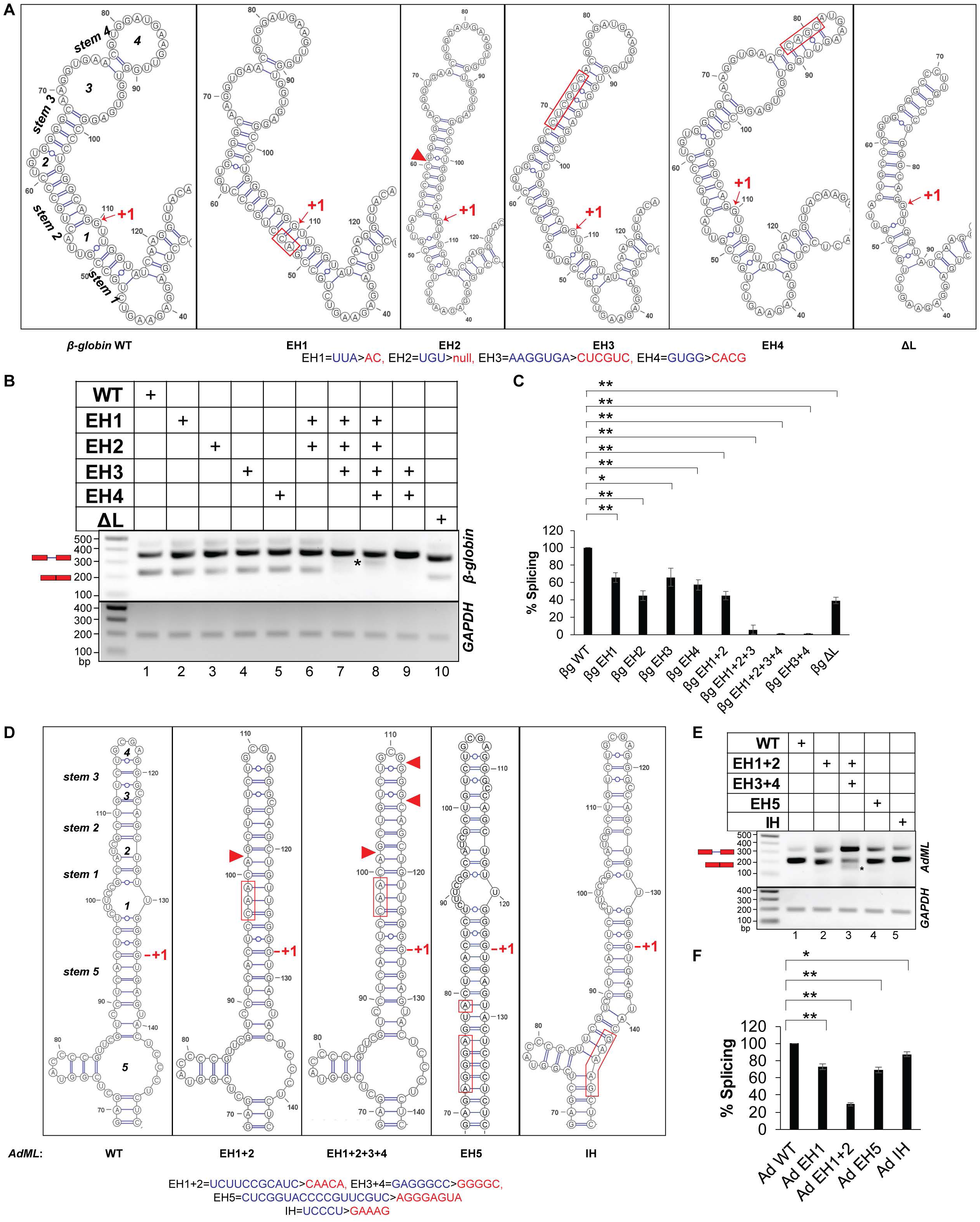
Secondary structure models and splicing activities of pre-mRNAs. (A) SHAPE-derived secondary-structure model of the WT *β-globin* segment immediately upstream of the 5′-SS (leftmost panel); the designation of loops and stems is noted; anticipated secondary structure models (based on the WT structure) of the exonic segment upstream of the 5′-SS of the mutants corresponding to hybridization/deletion of loops (EH/ΔL) are shown (remaining panels); substituted and deleted native sequence in the mutants are indicated with red rectangle and red triangle, respectively; the original and mutated sequences are shown at the bottom (also see Supplementary Figure S2A); the first nucleotide of the intron is marked with ‘+1’. (B) A representative image showing splicing products of WT, EH, and ΔL mutants of *β-globin* in transfection-based splicing assay; asterisks indicate aberrantly processed products; pre-mRNA and mRNA positions are marked with a red schematic; *GAPDH* amplification performed as an internal control shown at bottom. (C) Bar-graph showing changes in spliced mRNA products for mutants as a percentage of wild-type RNA and normalized with total RNA generated from the transfected plasmids (n = 3); error bars indicate standard deviation; statistical significance was tested to validate if EH/ΔL mutants of *β-globin* have lower splicing competence than the WT substrate; ‘*’ = 0.005<*p*≤0.05, ‘**’ = *p*≤0.005, N.S. = not significant. (D) SHAPE-derived secondary-structure model of WT *AdML* exonic segment immediately upstream of the 5′-SS (leftmost panel); anticipated secondary structure models of the exonic and intronic hybridization mutants (EH and IH) are shown; original and mutated sequences are shown at the bottom; also see Supplementary Figure S2B. (E) A representative image of splicing products of WT, EH, and IH mutants of *AdML* obtained by transfection-based splicing assay; asterisk and red schematic as in (B). (F) Bar-graph showing quantified mRNA along with the statistical significance of the differences; statistical significance was tested to validate if EH/IH mutants of *AdML* have lower splicing competence than the WT substrate.

A transfection-based assay indicated that splicing was significantly reduced in all EH mutants. While up to a 50% reduction was observed in individual EH mutants and the EH1+2 mutant, a complete or near complete abolition of splicing was observed in the other combinatorial mutants (EH1+2+3, EH1+2+3+4, EH3+4) (Figure 2B, 2C). We confirmed base-pairing between strands (i,e. elimination of the loops) in the *β-globin* EH3+4 mutant experimentally by *in vitro* SHAPE (Supplementary Figure S2C). In the mutant with all four terminal loops hybridized (EH1+2+3+4) (Figure 2A), authentic splicing was abolished, and an aberrant product was detected (Figure 2B, lane 8, aberrant product marked with an asterisk). The aberrant product matched the *β-globin* EH1+2+3+4 pre-mRNA with a deletion of a 63-nt region within the 5′ exon (between the 32^nd^ and 96^th^ nucleotides, highlighted by red borders in Supplementary Figure S3A, middle panel). We also generated a *β-globin* ΔL mutant in which loops 3 and 4 were replaced by a tetraloop (Figure 2A); this displayed a ∼50 % loss of splicing (Figure 2B, compare lanes 1 & 10, Figure 2C). In contrast to the EH mutants, the IH mutants of *β-globin* (Supplementary Figure S2A) showed up to an approximately two-fold enhancement of splicing (Supplementary Figures S2D, S2E).

We next examined if splicing was restored by reintroducing loop(s) into the 5′ exon of the *β-globin* EH1+2+3+4 mutant (Supplementary Figure S3A, middle panel). We added loops in the same general locations of Loops 1 and 2 in the WT substrate but composed of sequences different from WT, and called the new variant EH1+2+3+4+5L2 (Supplementary Figure S3A, right panel; the straight cyan line in the middle panel indicates the sequence of EH1+2+3+4 mutated to generate EH1+2+3+4+5L2). We observed that addition of the new loops eliminated the aberrant product of EH1+2+3+4 and slightly improved *bona fide* splicing. We also added a large loop in the general location of loops 3+4 of the WT substrate to generate a terminal loop (TL) in the EH1+2+3+4 mutant, and called it EH1+2+3+4+TL (Supplementary Figure S3A, left panel; the straight black line in the middle panel indicates the sequence of EH1+2+3+4 mutated to generate EH1+2+3+4+TL). Splicing in this variant was restored to ∼50% of WT levels (Supplementary Figure S3B, S3C).

Similar to the *β-globin* mutants, *AdML* EH mutants (Figure 2D) also displayed defects in splicing. The most severe defect was observed when all four loops upstream of the 5′-SS were hybridized (EH1+2+3+4); an aberrant product was also noticed (Figure 2E, compare lanes 1 & 3, 3F). Sequencing of this aberrantly processed product indicated that a GU dinucleotide (at positions 76 & 77, demarcated with red borders in Supplementary Figure S2B) can act as a cryptic 5′-SS and pair with the authentic 3′-SS. Hybridization-mutation at an intronic location of the 5′-SS-containing stem-loop (IH) (Figure 2D) produced a small (∼10%) defect in splicing (Figure 2E, compare lanes 1 & 5, Figure 2F). These results suggest that single-stranded regions immediately upstream of the 5′-SS may play essential roles in promoting splicing; however, these experiments have not clarified the role of specific nucleotide sequences within these exonic regions.

### Single-stranded sequence immediately upstream of the 5′-SS promotes splicing in a sequence- and context-specific manner

We assessed the importance of the nucleotide sequence within native exonic loops for splicing of constitutively spliced substrates. Here we focused on the loop regions of *β-globin* stem-loops 3 and 4. We made multiple purine-to-pyrimidine substitutions within the loops without disrupting the secondary structure of the stem-loop as predicted by ‘RNAstructure’ (33). Among these mutants, only mutant Mt (see Figure 3A for mutation, 3B and 3C for predicted secondary structure models of Mt and WT) exhibited a ∼20 % reduction in splicing efficiency compared to the WT (Figure 3D and 3E). The remaining mutants did not exhibit any change in splicing. Due to high degeneracy in consensus sequence for binding of splicing factors and the presence of numerous splicing factors inside the nuclei with varied sequence specificity, mutations without an effect on splicing is not useful for evaluating the importance of an RNA sequence on splicing. Therefore, the results of the mutants other than that of Mt are not shown.

**Figure 3.**
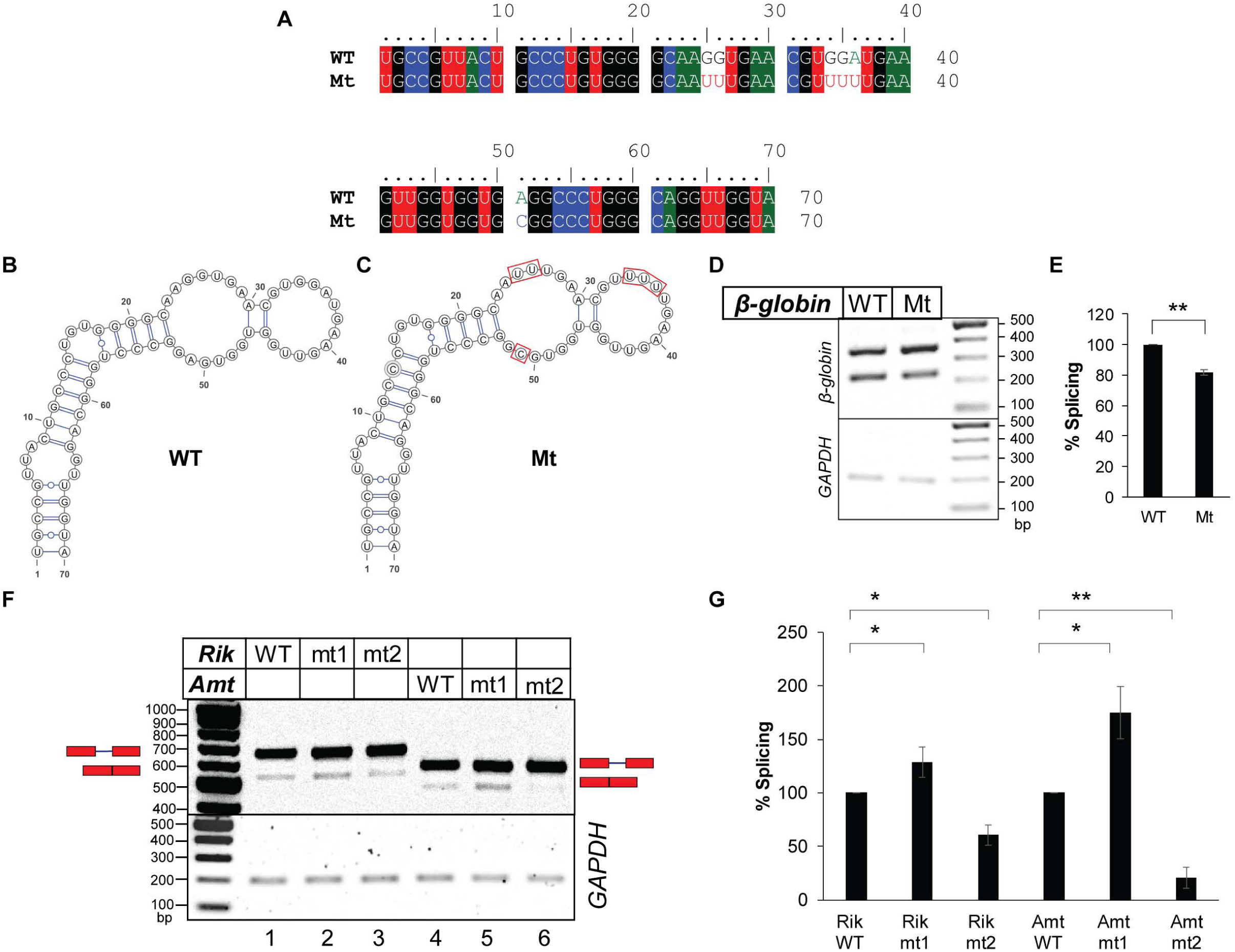
Sequence- and context-specific effects of single-stranded nucleotide sequence in the exonic segments immediately upstream of the introns on splicing. (A) Sequence alignment of the 5′ exonic segment of WT and mutant *β-globin* variants. (B & C) Secondary structure models of the above exonic segment of WT *β-globin* (B) and the mutant *β-globin* (C) (both predicted by ‘RNAstructure’), which are identical to experimentally derived secondary structure of WT *β-globin* shown in Supplementary Figure S2A. (D) Transfection-based splicing assay of the WT and the mutant *β-globin* variants. (E) Plots of quantified band-intensities shown in (D) (n=3); statistical significance was tested to validate if the splicing competence of Mt mutant of *β-globin* is less than that of WT *β-globin*; ‘*’ = 0.005<*p*≤0.05, ‘**’ = *p*≤0.005, N.S. = not significant. (F) Splicing products of mouse pre-mRNAs *2610507B11Rik* IVS17 (*Rik*) and *Amt* IVS1 (*Amt*) and their mt1 and mt2 mutants (see Supplementary Figure S5) visualized in agarose gel (representative); positions of pre-mRNAs and mRNAs are marked and *GAPDH* amplification used as an internal control shown. (G) A bar-graph showing comparison of spliced mRNAs normalized with total RNA generated from the transfected plasmids; error bars (n=3) indicate standard deviation; statistical significance was tested to validate if mt1 mutants have higher and mt2 mutants have lower splicing competence than the WT substrate; ‘*’ = 0.005<*p*≤0.05, ‘**’ = *p*≤0.005.

Next, we examined if expansion of single-stranded segments (by disrupting base-pairing in the stems) immediately upstream of the 5′-SS of *β-globin* and *AdML* enhances splicing. We generated exonic de-hybridization (ED) mutants by removing base pairs from stems 1, 2, 3, or 4 of *β-globin* (Supplementary Figure S4A) and stems 1, 2, 3, or 5 of *AdML* (Supplementary Figure S4D). Splicing of these substrates revealed no clear trend. Only ED1 of *β-globin* exhibited splicing enhancement while the rest did not. On the other hand, all but one *AdML* ED mutant exhibited a reduction in splicing; only (ED1+5) exhibited no significant change. (Supplementary Figure S4B, S4C, S4E, S4F). Since the base substitutions to generate ED mutants were random (without any emphasis on specific sequence), the unpredictable splicing changes observed were likely due to random changes in the strength of available ESE and ESS elements of *β-globin*.

Next, we examined how replacing a single-stranded as well as a base-paired exonic sequence immediately upstream of the 5′-SS with a strong single-stranded heterologous hexameric ESE (4) altered the splicing activity of two mouse pre-mRNA substrates exhibiting intron retention. The mouse pre-mRNAs chosen have sufficient reads mapped to the retained introns in both *in vivo* and *in vitro* datasets (*2610507B11Rik* IVS17 and *Amt* IVS1, Supplementary Figure S5A). Secondary structure models based on their *in vitro* icSHAPE reactivities were generated using ‘RNAstructure’ (33) (Supplementary Figures S5B and S5C). Mutant ‘mt1’ was generated by replacing a base-paired native sequence with a single-stranded, high-ranking ESE (UCAUCG-ranked 35) (4) and ‘mt2’ was generated by replacing a loop sequence with another strong ESE (GAAGAA-ranked 17). *Rik* IVS17 mt1 exhibited a small increase in splicing efficiency (Figure 3F, 3G); on the other hand, *Rik* mt2 exhibited a decrease in splicing efficiency. mt1 of *Amt* IVS1 was created by replacing the base-paired native sequence with AAGAAA (ESE rank 25), and in mt2 of *Amt* IVS1 the native sequence of the loop was replaced with AAGAAC (ESE rank 102). *Amt* IVS1 mt1 exhibited an increase in splicing while mt2 exhibited a decrease (Figure 3F, 3G).

Overall, these results suggest that insufficient single-stranded ESE sequences within exons immediately upstream of the retained introns could be responsible for splicing defects, and that loops could display context-specific ESE which may not be replaceable with a heterologous ESE. Specific unwinding of the naturally hybridized exonic sequence immediately upstream of the retained introns and demonstration of its functional relevance in promoting splicing is difficult. However, the observation of enhanced splicing in the mt1 mutants of both mouse pre-mRNA substrates suggests that a single-stranded segment in place of the base-paired stem in the exonic region immediately upstream of the 5′-SS could promote splicing of the retained introns in a sequence/context-specific manner.

### Exonic base-pairs immediately upstream of the retained introns appear to occlude ESEs transcriptome-wide

The results shown above suggest that pre-mRNA-specific ESEs are present in the single-stranded loop regions and may be occluded within the base-paired regions immediately upstream of the mouse retained introns. We therefore asked if this base-pairing occludes ESE sequences throughout the transcriptome. We chose the set of hexameric ESE, ESS, and neutral sequences identified by Ke *et al* (4) and examined their occurrences in the exonic segments upstream and downstream of both retained and spliced introns across the transcriptome. Although the strength of ESE and ESS sequences is context-dependent, the observation of any distinct transcriptome-wide trend will likely provide a general idea regarding the distribution of ESE and ESS sequences with respect to the exonic secondary structure. A 70-nt stretch upstream or downstream of the intron was considered the ‘target region’, and the next 80-nt long exonic segment upstream or downstream was regarded as the ‘non-target region’. Figures 4A and 4B show combination (violin + box) plots of distribution of recognized hexameric ESE and ESS, as well as neutral hexameric sequences (4) in both upstream (5′) and downstream (3′) exonic segments of retained and spliced introns in the *in vivo* dataset. Higher ESE and lower ESS sequence counts were observed in both 5′ and 3′ exonic ‘target regions’ flanking retained introns than were observed in similar regions flanking spliced introns (Figure 4A and 4B). ‘Non-target regions’ in the 3′ exons but not the 5′ exons showed a similar difference.

**Figure 4.**
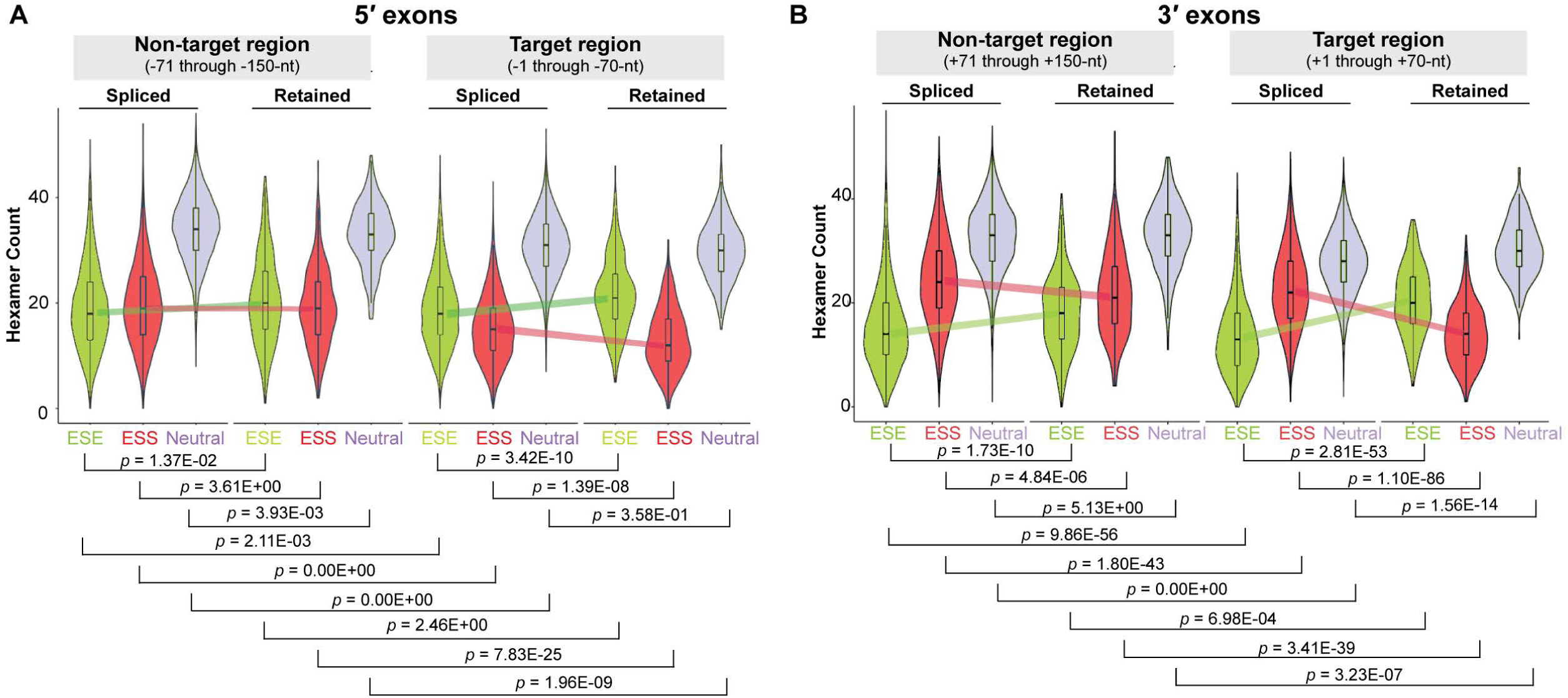
Distribution of ESE/ESS counts in the exonic segments flanking the retained introns. (A) Counts of occurrences of hexameric ESE, ESS, and neutral sequences (green, red, and blue, respectively) (4) in the -1 to -70-nt (target region) and -71 to -150-nt (non-target region) ranges upstream of the 5′-SS of retained and spliced introns (B) Similar counts in the +1 to +70-nt (target region) and +71 to +150-nt (non-target region) ranges downstream of the 3′-SS of retained and spliced introns; *p*-values are indicated; colored lines indicate deviation of medians between spliced and retained introns.

In order to gain an idea of the average splicing potential for specific exonic segments (both target and non-target regions), we calculated their overall ‘splice score’ by summing up individual ESE and ESS scores. The values for hexameric ESE and ESS sequences (positive and negative, respectively) were obtained from Ke, *et al.* (4) (and were therefore calculated based on the potential of these sequences to regulate splicing within the context of their experiments). We observed ‘splice scores’ of target regions flanking the retained introns to be higher than regions flanking the spliced introns (Supplementary Figure S6A, S6B). Also, in the case of retained introns, the splice scores for target regions were higher than those for non-target regions.

Since the presence of common motifs within the exonic segments flanking the retained introns might indicate similarity in their splicing regulation, we looked for *de novo* motifs in exonic segments upstream of retained introns in the *in vivo* dataset versus the control group (spliced introns) using MEME (49) and MAST (50). The sequence motif CTTCTG and its reverse complement (sequence logos shown in Supplementary Figure S6C) occurred 122 and 235 times respectively within the exonic segments upstream of 207 retained introns. The CTTCTG hexamer as well as its reverse complement were classified as ESEs in a previous study (4).

Since occlusion of ESE sequences by base-pairing potentially inhibits their ability to promote splicing (27,51), and the exonic segments flanking the retained introns are on average enriched in hexameric ESE sequences identified by Ke et al (4), we hypothesized that occlusion of ESE or ESE-like sequences present in the 5′ exonic region by means of base-pairing could be one of the reasons for intron retention. However, comparison of icSHAPE reactivity (Figure 1B, C) and ESE count (Figure 4A, 4B) between retained and spliced introns indicated a more significant icSHAPE reactivity difference and a less significant ESE count difference in the ‘target region’ of upstream exons compared to those of downstream exons. These findings suggest that base-pairing of the ‘target regions’ may have additional purposes beyond regulating the exposure of ESEs.

### WT *β-globin* exhibits greater degree of single-strandedness than its EH3+4 mutant *in vivo*

The hybridization mutations are planned based on ‘static’ *in vitro* models of full-length pre-mRNAs. Therefore, the questions remain whether the mutants acquire the expected structure upon mutagenesis *in vivo*. In order to explore possibilities of how the exonic loops in the target region promote splicing *in vivo*, we wanted to examine the secondary structural features of *β-globin* WT and the EH3+4 mutant *in vivo*. We carried out *in vivo* DMS-footprinting of the 5′ exon-intron junction region of *β-globin* variants expressed from the transfected minigene. *In vivo* DMS-reactivity is shown in Figure 5. The DMS reactivity of A and C residues are mapped onto the SHAPE-derived secondary structure model of *β-globin* (Supplementary Figure S7). It is anticipated that an ensemble of structural states during the course of splicing are represented *in vivo.* The exonic region corresponding to stem-loops 3 and 4 in the splicing-competent WT substrate showed higher reactivity than that in the splicing-defective EH3+4 mutant substrate. Significantly, we also observed higher DMS-reactivity at a number of additional segments in both exons and introns outside of stem-loops 3 and 4 in the WT substrate compared to the EH3+4 mutant. In summary, *in vivo* WT *β-globin* exhibited more single-strandedness within as well as beyond the exonic segment immediately upstream of the 5′-SS compared to the EH3+4 mutant. Since protein-coding RNAs tend to be less structured *in vivo* than *in vitro* due to structural modulation predominantly caused by protein binding (15,52), we surmise that the exonic loops promote recruitment of splicing factors that in turn may induce structural modulation of the pre-mRNA.

**Figure 5.**
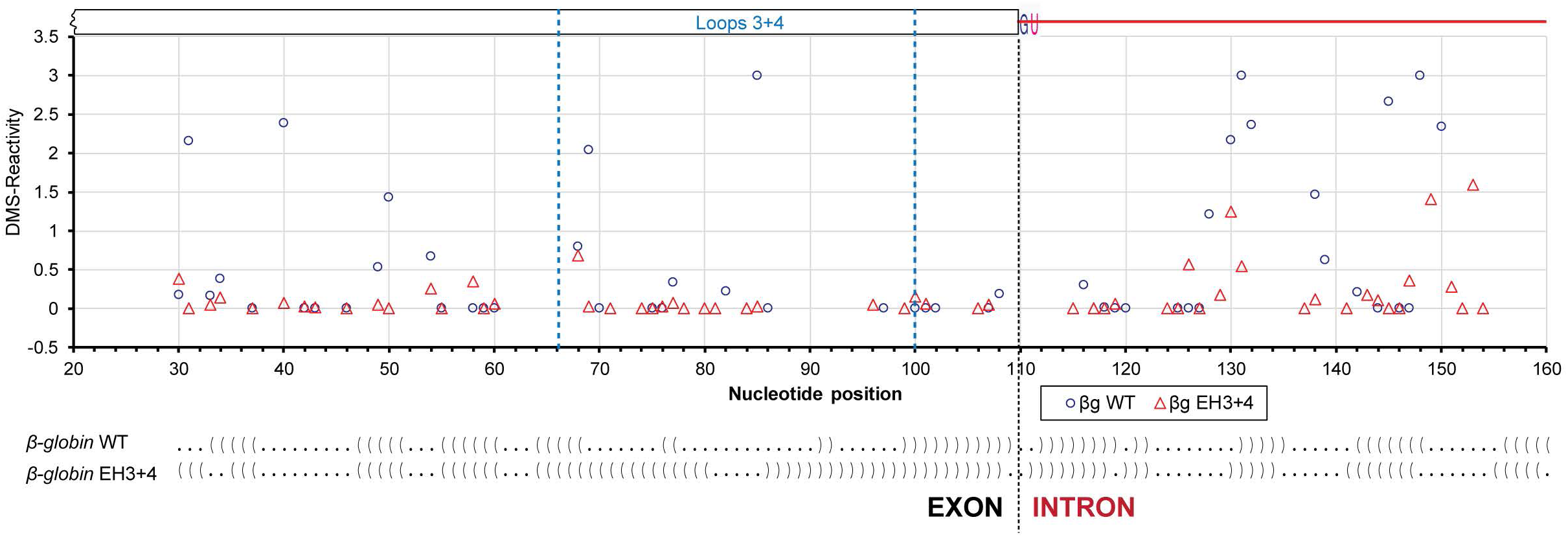
*In vivo* structural probing of *β-globin* WT and its EH3+4 mutant expressed from transfected minigenes. Quantified reverse transcription stops (DMS-reactivity) of individual nucleotides (A & C only) of *β-globin* WT (o) and EH3+4 mutant (Δ) are plotted against nucleotide positions; dot-bracket notations, which are aligned with the nucleotide positions in the plot, of the SHAPE-derived secondary structure models of protein-free WT *β-globin* and its EH3+4 mutant are shown at the bottom (dot = unpaired nucleotide, bracket = base-paired nucleotide).

### Single-strandedness immediately upstream of the intron is important for structural remodeling of *β-globin* and *AdML* by SR proteins

The results described above led us to examine if binding of splicing factors causes structural modulations *in vitro*, and whether such remodeling is regulated by secondary structure of the exonic segment immediately upstream of the 5′-SS. We focused on SR protein-mediated structural modulations of the model substrates, *β-globin* and *AdML* since they are known to be activated for splicing by the SR proteins (53-55).

We examined data from multiple transcriptome-wide crosslinking and immunoprecipitation experiments followed by RNA-seq (CLIP-seq), which identified that SR proteins bind predominantly to exonic regions within ∼50-nt of the 5’-SS and 3′-SS (8-10). Enhanced CLIP (eCLIP) data (9) for binding of three different SR proteins (SRSF1, SRSF7, and SRSF9) to *SRSF1* exon 1 are shown in Figure 6A (www.encodeproject.org), which shows the presence of eCLIP peaks for all three SR proteins within the ∼70-nt region immediately upstream of *SRSF1* intron 1. We used EMSA to assess binding of SR protein SRSF1 to *β-globin* and *AdML* pre-mRNAs (500 nM unlabeled pre-mRNAs with radiolabeled tracer) (Supplementary Figures S8A, S8B, S8C). The stoichiometric binding assay revealed that ∼30 molecules of SRSF1 are needed to saturate a WT *β-globin* pre-mRNA molecule (Supplementary Figure S8A, lane 6) and ∼9 molecules for a WT *AdML* molecule (Supplementary Figure S8B, lane 6). This difference may be reflective of the much longer 3′ exon of *β-globin* (200-nt) compared to that of *AdML* (50-nt). A similar experiment with SRSF2 indicated that 10 molecules are required for saturation of a *β-globin* molecule (Supplementary Figure S8C, lane 5). Overall, these studies show binding of multiple molecules of SR proteins to a pre-mRNA molecule, which may be related to dose-dependent splicing regulation by SR proteins (56).

**Figure 6.**
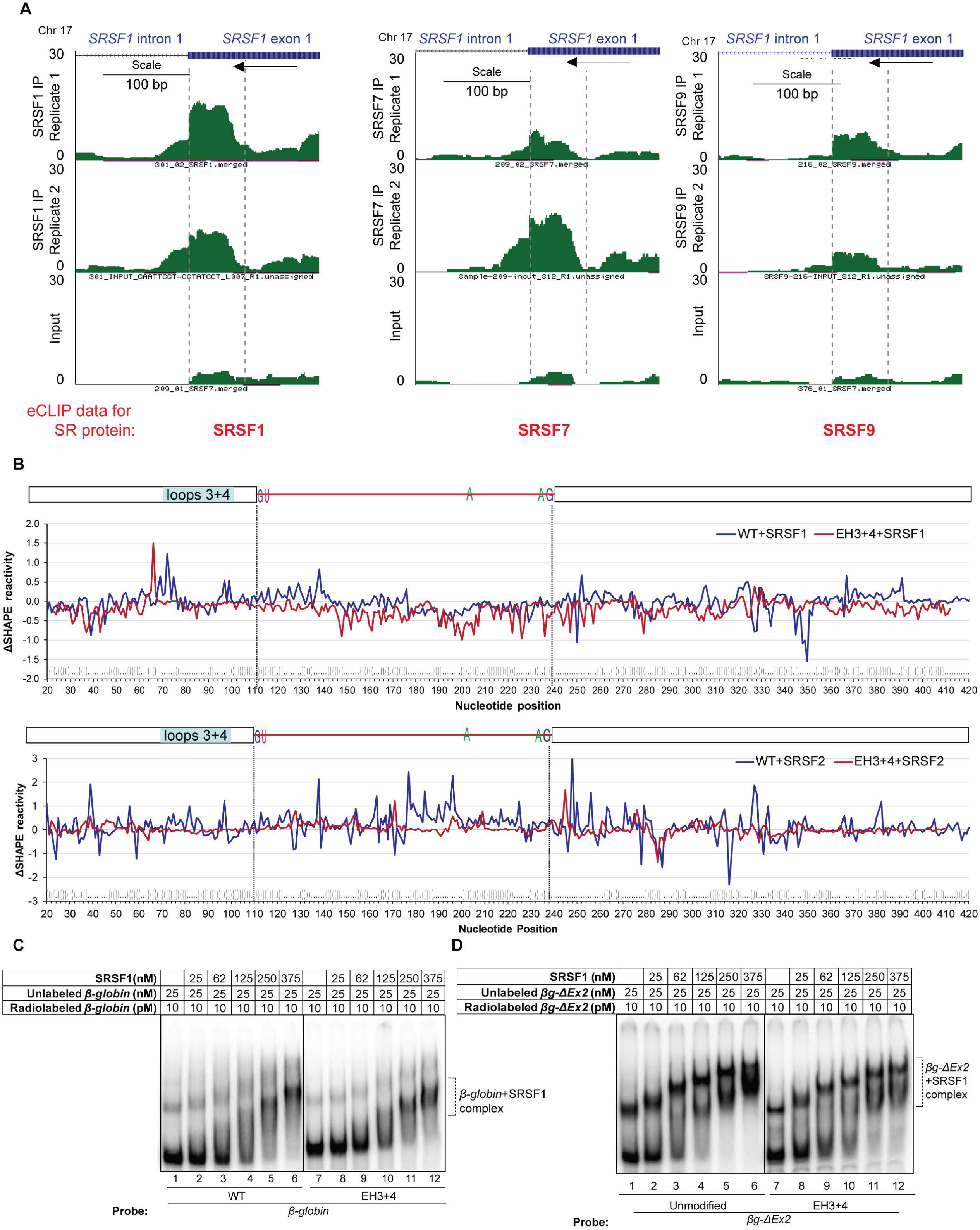
Effect of exonic loops immediately upstream of the 5′-SS to functional SR protein binding and pre-mRNA structural modulation. (A) Genome browser views of 5′ exon of *SRSF1* IVS1 with mapped SR protein (SRSF1, SRSF7, and SRSF9) eCLIP reads (9) showing the exonic region immediately upstream of the 5′-SS to overlap with high confidence SR protein binding sites; mapped SR protein read densities from two eCLIP biological replicate experiments performed in HepG2 cells and one size-matched input are plotted at the same *y*-axis scale; scale bar and the directions of transcription (by an arrow) are denoted (www.encodeproject.org); 70-nt exonic segment immediately upstream of the intron is marked with two vertical broken lines. (B) (Top) SHAPE reactivity differential (ΔSHAPE reactivity) of free and SRSF1-bound *β-globin* WT (blue line) and EH3+4 mutant (red line). (Bottom) SHAPE reactivity differential of free and SRSF2-bound *β-globin* WT (blue line) and EH3+4 mutant (red line); dot-bracket notation of WT *β-globin* secondary structure is shown at the bottom, which is aligned with nucleotide positions; position of 70-nt at the 3′ end of the upstream exon (−1 through -70-nt) is indicated. (C) Comparison of SRSF1 binding to full-length WT *β-globin* and its EH3+4 mutant by EMSA. (D) Comparison of SRSF1 binding to *βg-ΔEx2* variants (truncated *β-globin* lacking the downstream exon) with and without EH3+4 mutation (compare lanes 4, 5, and 6 with 10, 11, and 12, respectively).

We next probed structural modulation of WT *β-globin* and its EH3+4 mutant upon binding of SRSF1 and SRSF2, and of WT *AdML* upon binding of SRSF1 by monitoring changes to *in vitro* SHAPE reactivity. WT *β-globin* and its EH3+4 mutant were saturated independently with SRSF1 and SRSF2, and WT *AdML* with SRSF1. The resulting SHAPE reactivity differentials (ΔSHAPE-reactivity) were plotted against individual nucleotide positions (see Figure 6B top panel for *β-globin*+SRSF1, Figure 6B bottom panel for *β-globin*+SRSF2, Supplementary Figure S8D for *AdML*+SRSF1). Positive and negative ΔSHAPE-reactivity values indicate gain and loss of reactivity upon SR protein binding and imply an increase and decrease of nucleotide flexibility respectively. ΔSHAPE-reactivities (both loss and gain) of the *β-globin* EH3+4 mutant were diminished compared to the reactivities of the WT substrate. We also analyzed the raw SHAPE data of *β-globin* and *β-globin*+SRSF1 by 90% Winsorization and compared the resulting ΔSHAPE-reactivity with that described above obtained by ‘trimming’. Both methods yielded comparable results (*p* = 1.53E-02).

As performed for the retained and spliced introns, we calculated the PU values for hexamers of *β-globin* and *AdML* using MEMERIS, RNAplfold (L=70, W=120) and LocalFold (L=70, W=120). RNAplfold and LocalFold values were almost identical while MEMERIS values were somewhat different (Supplementary Figure S9). No obvious positive correlation between the SHAPE reactivity of protein-free pre-mRNAs as well as SRSF1-bound pre-mRNAs and the PU values for different segments of the model substrates were observed (Supplementary Table S2). Similarities in trend of the curves of SHAPE reactivity of protein-free as well as SRSF1-bound pre-mRNAs and the PU values in certain small patches were observed by visual inspection of the plots (Supplementary Figure S9). PU values obtained by both MEMERIS and LocalFold of the PPT region (215-239-nt) of *β-globin* are distinctly high but the protein-free pre-mRNA shows only eight unpaired nucleotides in this region (Supplementary Figure S2A). Interestingly, SR protein binding to the WT substrates appeared to maintain, enhance, or weakly suppress the flexibility of the PPT region while SR protein binding to the EH3+4 mutant appeared to strongly suppress the flexibility of this region (Figure 6B): the *p*-value of the difference in ΔSHAPE reactivity in the PPT and its flanking areas [150-250 nt] of *β-globin* WT+SRSF1 and *β-globin* EH3+4+SRSF1 mutant is 3.90E-08; and that of *β-globin* WT+SRSF2 and *β-globin* EH3+4+SRSF2 is 1.88E-09. This might be reflective of the requirement of a more single-stranded PPT region for splicing (57).

We further compared SRSF1 binding to WT and EH3+4 mutant of *β-globin* by EMSA (Figure 6C). No difference in SRSF1 binding efficiency between the WT and the mutant *β-globin* was clearly discernible, except that the band formed upon addition of 375 nM SRSF1 was slightly less compact in the case of *β-globin* EH3+4 than in WT *β-globin*. Since the presence of a long (200-nt) 3′ exon could suppress any binding difference, we generated a *βg-*ΔEx2 construct with the 3′ exon of *β-globin* removed. The band formed in the presence of 375 nM SRSF1 with 25 nM *βg-* ΔEx2 is clearly more compact than the one formed with 25 nM *βg-*ΔEx2-EH3+4 (Figure 6D, compare lanes 6 with 12). These results suggest that loops in the exonic segments immediately upstream of the 5′-SS govern the nature of SRSF1 binding to *β-globin*.

### Single-strandedness immediately upstream of the intron is important for E-complex assembly

To assess the effect of exonic secondary structure on formation of the splicing complex, we examined assembly of the E-complex on *β-globin* WT and its EH3+4 and EH1+2+3+4 mutants in nuclear extract. Native sequence pre-mRNA formed the H-complex at 0 min (i.e., upon mixing) and E-complex in 15 min (Figure 7A, lanes 1 & 2). E-complex formation was compromised in both with EH3+4 and EH1+2+3+4 mutants (Figure 7A, lanes 13-18), in solid agreement with the results from the *in vivo* splicing assay results in Figure 2. Additionally, we observed that pre-incubation of native *β-globin* (performed with ∼10 pM radiolabeled pre-mRNA) with SRSF1 resulted in greater efficiency of E-complex formation (Figure 7A, compare lanes 2 & 8). Pre-incubated substrate was able to survive a challenge of ∼1000-fold excess (unlabeled) substrate while untreated substrate was not (compare lanes 3 & 9). As expected, *β-globin* of native sequence was able to form spliceosomal complexes A and B, but its EH3+4 mutant was not (Figure 7B). A schematic of this function of exonic secondary structure is presented in Figure 7C, D.

**Figure 7.**
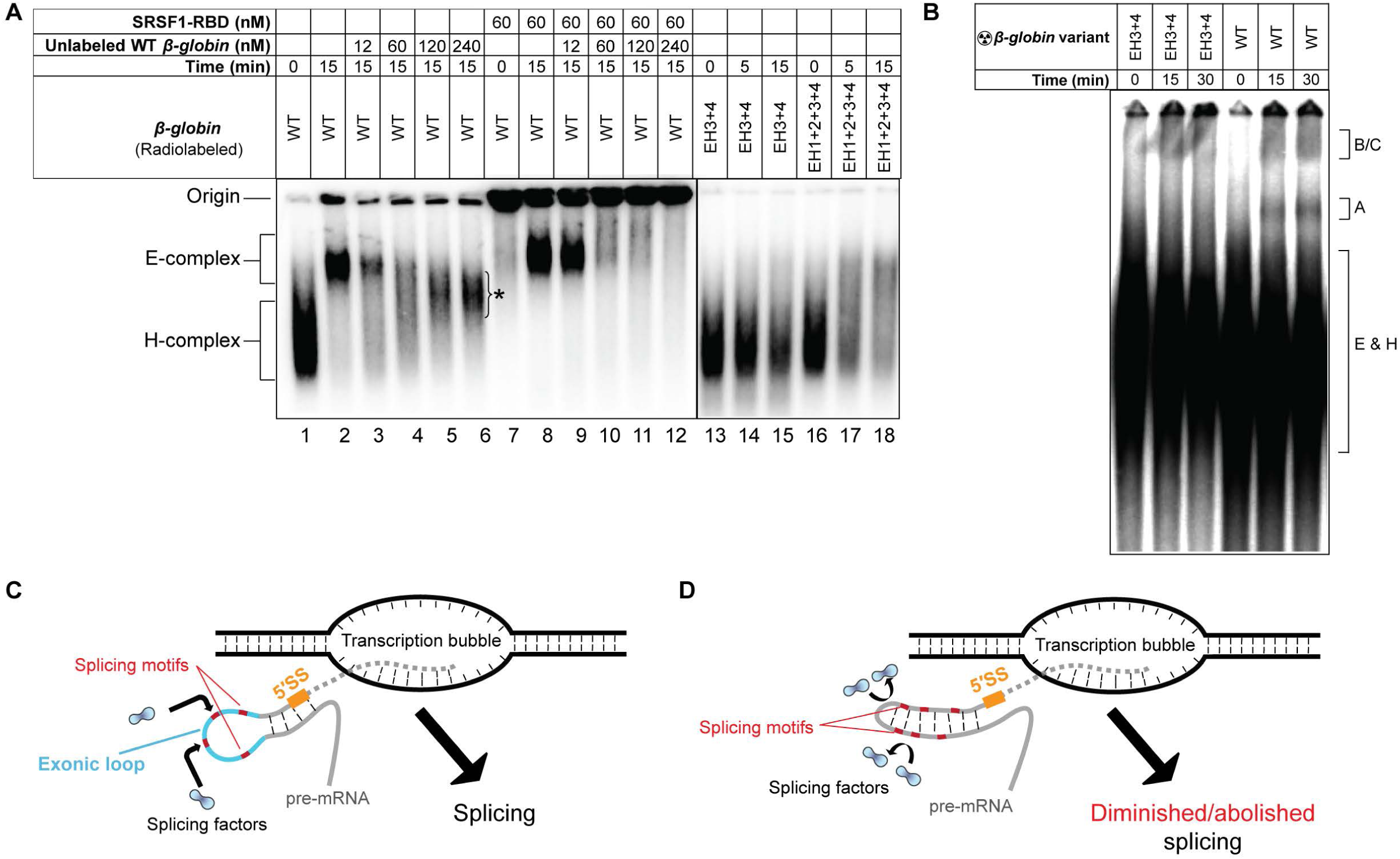
Importance of exonic loops immediately upstream of the 5′-SS for assembly of the early spliceosome. (A) Comparison of E-complex assembly with 10 pM radiolabeled *β-globin* pre-incubated with SRSF1 in the presence or the absence of 12 nM untreated unlabeled competitor *β-globin* (compare lanes 3 and 9); E-complex assembly assay performed with *β-globin* EH3+4 and EH1+2+3+4 mutants (lanes 13-18); an asterisk indicates undefined pre-mRNA complexes; nuclear extract from the same preparation is used in all experiments. (B) Spliceosomal complex (A, B/C) assembly assay with *β-globin* WT and its EH3+4 mutant. (C & D) Schematics depicting the role of exonic secondary structure in spliceosome assembly; a pre-mRNA in the process of being released from the transcription bubble is shown; solid grey lines indicate exons and dotted grey lines intron; orange line indicates 5′-SS, cyan line the exonic loop(s) immediately upstream of the 5′-SS, and short red lines putative splicing motifs. In case of splicing competent introns, the exonic segment immediately upstream of the 5′-SS contain single-stranded segments, which promote recruitment of appropriate splicing factors for splicing. Stems in place of exonic loops either occlude the splicing motifs within or provide steric hindrance to recruitment of a splicing factor at nearby sites or both, thus diminishing splicing.

## DISCUSSION

Exonic stem-loops are postulated to provide structural context to splice sites, predominantly by regulating exposure of ESE and ESS sequences (27,58) and influencing their ability to recruit splicing factors. We observed that exonic segments immediately upstream of retained introns transcriptome-wide are extensively base-paired and enriched in at least one set of previously identified hexameric ESE sequences compared to those upstream of the spliced introns. In contrast, the downstream exonic segments, while also enriched in ESE sequences compared to their spliced counterparts, appeared to be more single-stranded. With select pre-mRNA substrates, we observed that loops present in the upstream exonic segments promote splicing in a sequence- and context-dependent manner. *In vitro*, we observed that these loops facilitate recruitment of SR proteins and induce structural modulation of the pre-mRNA, perhaps exposing additional splice signals. *In vivo*, the pre-mRNA appeared more single-stranded in the presence of the loops than in their absence, suggesting involvement of splicing factors in structural modulation of the pre-mRNA. In addition to being enriched in ESE sequences, the exonic segments flanking the retained introns also exhibited a dearth of ESS sequences identified by Ke et al (4) compared to the exonic segments flanking the spliced introns. This might suggest an apparent higher splicing potential of the exonic segments flanking the retained introns than those of the spliced introns. Based on these observations, we propose the following model for mammalian splicing regulation: Single-stranded ESEs immediately up- and down-stream of splice sites bind sequence-specific protein factors which modulate the pre-mRNA structure to sequentially recruit components of the spliceosome core. Extensive base-pairing immediately upstream of the retained introns reduces functional interactions between ESEs and splicing factors. Exposed ESEs immediately downstream of the retained introns cannot sufficiently substitute for the function of occluded ESEs in the 5′-exon. Promotion of splicing triggered by unwinding of the exonic segments upstream of the retained introns likely is not interfered by exposed ESS sequences due to their low counts. A previous study suggested that ∼40-nt exonic regions immediately upstream but not downstream of the retained introns are highly base-paired in *Arabidopsis thaliana* (14), suggesting that a similar mechanism of intron retention might also prevail in higher plants.

It will be intriguing to determine if the structural feature(s) associated with the exonic segments are correlated with alternative, biologically significant intron retention events. However, in the absence of structural data (e.g. icSHAPE reactivity) from multiple cell lines or cells under different conditions, we can only hypothesize about the possibility of exonic secondary structure contributing to alternative retention of an intron in different circumstances. Biologically significant alternative intron retention regulated by exonic secondary structure might be effectuated by different RNA polymerase II elongation rates affecting co-transcriptional pre-mRNA folding, differential activities of helicases, or both either in different cell-types or in the same cell-type under different conditions. Differential folding of a pre-mRNA may not only change the accessibility of specific sequences within the exons but could also affect the efficiency of unwinding by helicases, which in turn may depend on the strength of base-pairing within the stem.

Observation of minor correlation between PU values obtained by various methods and SHAPE/icSHAPE reactivity in a few segments/patches could stem from our incomplete understanding of the principles of RNA folding. The prediction of local structure is dependent on fixed ‘motif-length’ and fixed L and/or W parameters and thus, could be limited. Furthermore, *in vivo*, RNA could fold differently in the presence of interacting proteins and RNA folding is often regulated by the transcription elongation rate; these possibilities cannot be easily considered for calculation of PU values. Overall, our data indicates a lack of a strong obvious correlation between PU values and SHAPE/icSHAPE reactivity and emphasize the need for experimentally determined RNA structure for a clearer understanding of RNA structure-function relationship.

There are many additional questions that need to be answered to better appreciate the impact of exonic stem-loops in splicing. Splicing efficiency of full-length pre-mRNAs with multiple exons and introns could depend upon a number of factors including *cis*-acting signals present at any distance from individual introns (59). It is not yet clear how these signals integrate with exonic stem-loop properties to regulate splicing. It is also not known if the binding of other splicing factors besides SR proteins is regulated by exonic secondary structure. Additionally, our data does not reflect how the extent of single-strandedness in the upstream exonic segment correlates with splicing efficiency. For example, *AdML* splices more efficiently than *β-globin* in spite of the fact that the former contains a less single-stranded region immediately upstream of the intron than the latter.

In this report, we experimentally verified the effects of base-pairing and single-strandedness of the exonic segment on splicing using pre-mRNAs containing short introns. It is not clear if such a regulatory mechanism is universal for all pre-mRNAs. Further studies exploring a genome-wide secondary structural profile of pre-mRNAs in cells with arrested splicing will reveal the extent to which single-stranded segments upstream of the 5′-SS are required for splicing. Additionally, our data show that SR protein-mediated structural modulation of the pre-mRNA is important for splicing but it does not explain whether or how variation in intron retention (often achieved by varying the expression level of SR proteins (8)), is regulated by SR protein-mediated pre-mRNA structural remodeling. The mechanism behind the compensatory effects of multiple SR proteins on splicing (8) and the likely link to pre-mRNA structural modulation are also unclear and requires further investigation. Finally, as observed with exonic secondary structure, the intronic secondary structure may also have a regulatory role in splicing. For example, certain types of intronic hybridization have been proposed to be involved in the promotion of splicing (60). Our preliminary observations of intronic hybridization (IH) mutants of *β-globin* showing distinct enhancement in splicing and the IH mutant of *AdML* exhibiting a small defect in splicing suggest that intronic secondary structure could have contributed to splicing regulation. However, regulatory schemes mediated by intronic secondary structure is likely different and different sets of splicing factors could be involved. Further studies are required to unravel the role of intronic secondary structure on splicing.

## AUTHOR CONTRIBUTION

KS designed and performed majority of the experiments, analyzed the data, and wrote the paper. TB analyzed data and wrote the paper. MF performed protein purification, cloning, and site-directed mutagenesis experiments, and helped writing the paper. RS and WE performed and analyzed the icSHAPE data and wrote the paper. GG planned experiments, analyzed data, wrote the paper, and supervised the project.

## ACKNOWLEDGMENT

The authors acknowledge Drs. Stephan Leung and Suhyung Cho for experiments at the early stage of this work, Drs. Gene Yeo and Stefan Aigner for providing expertise on eCLIP data analysis, and Drs. Matt Daugherty and Shankar Subramaniam for critical discussion on bioinformatic analyses.

## Supplementary Figure File

**Supplementary Figure S1.**
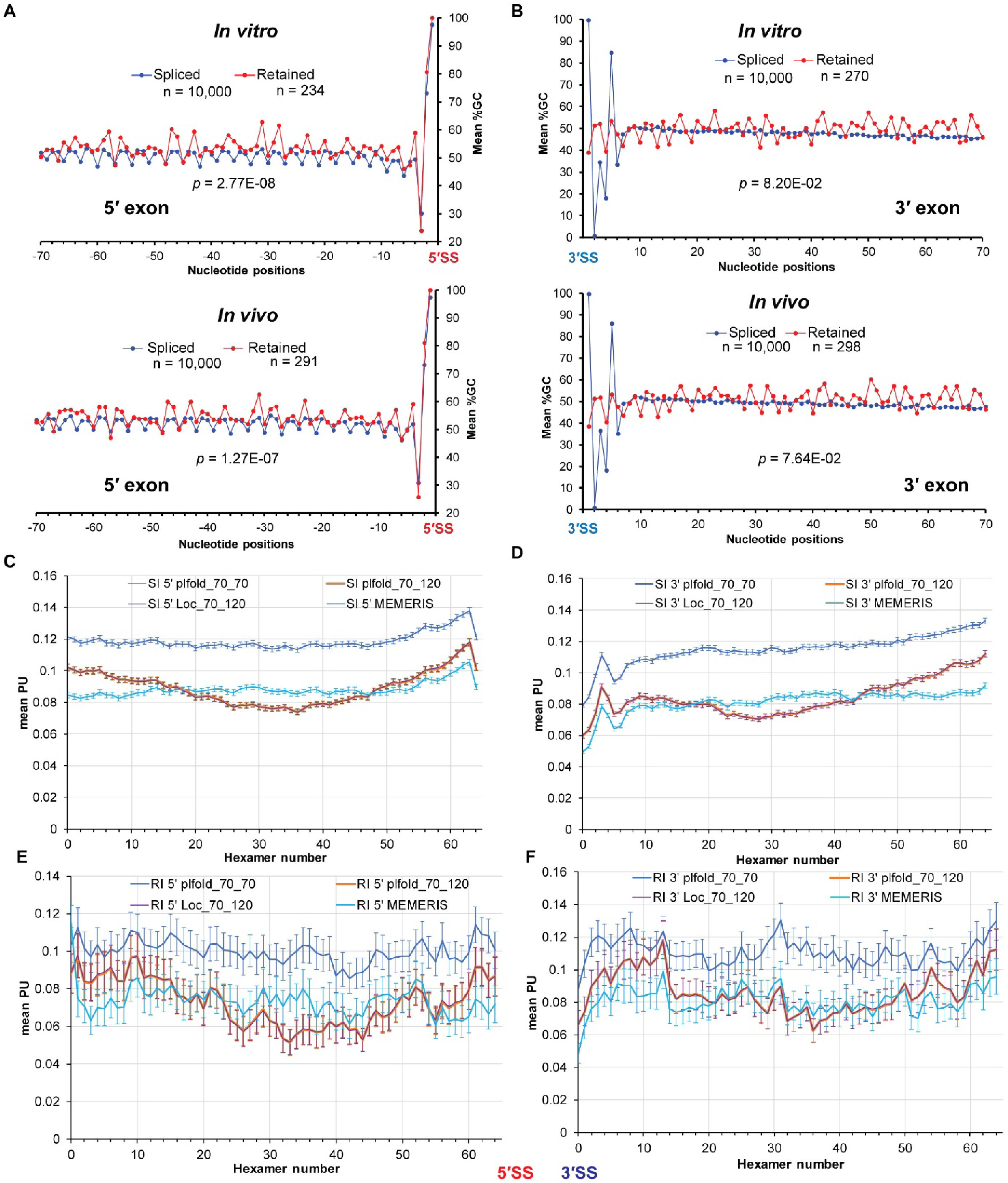
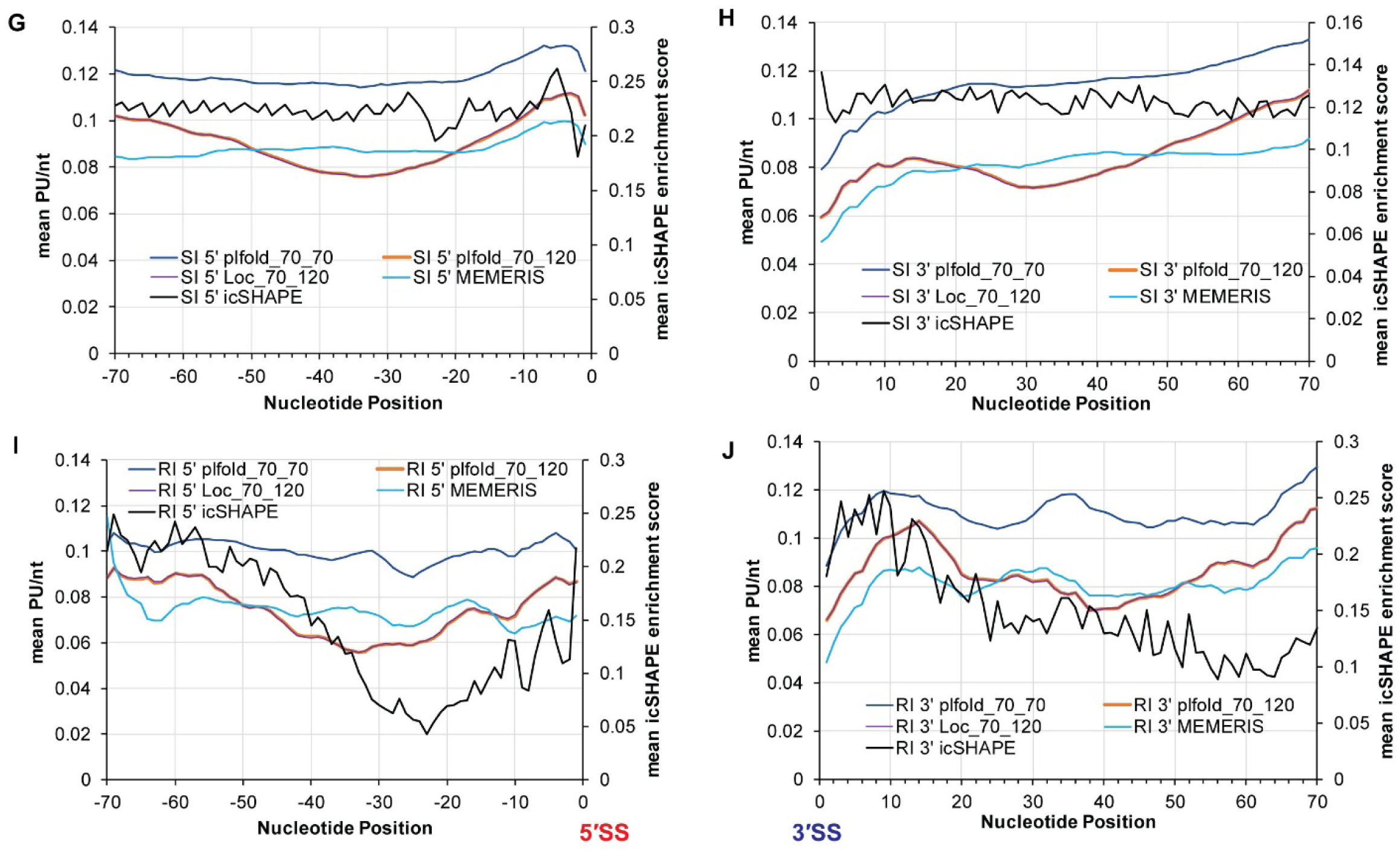
Comparison of properties of exonic segments flanking the retained and the spliced introns. (A) Mean G/C content of the exonic segments upstream of retained (red line) and spliced (blue line) introns in the *in vitro* (top) and *in vivo* (bottom) datasets; *p*-value of the difference in G/C contents is indicated; nucleotide positions upstream of the intron (negative digits starting from the 5′-SS) are marked. (B) Mean G/C content of the downstream exonic segments in *in vitro* (top) and *in vivo* (bottom) datasets. (C, D, E, F) Mean ‘probability of being unpaired’ or PU values of the hexamers of the exonic segments upstream (C) and downstream (D) of the spliced introns (SI) and upstream (E) and downstream (F) of the retained introns (RI) calculated by LocalFold with L=70, W=120 (Loc_70_120), MEMERIS, RNAplfold with L=W=70 (plfold_70_70), and RNAplfold with L=70, W=120 (plfold_70_120) (plfold_70_120 curve almost overlaps with Loc_70_120 curve); error bars indicate standard error of the mean; consecutive hexamer positions (0 – 64) from 5′ to 3′ end of the 70-nt long ‘target region’ are indicated along *x*-axis. (G, H, I, J) Comparison of mean icSHAPE enrichment score and mean PU values *per nucleotide* of the exonic segments calculated by LocalFold with L=70, W=120, MEMERIS, RNAplfold with L=W=70, and RNAplfold with L=70, W=120 flanking the 5′SS (G) and 3′SS (H) of the spliced introns (SI), 5′SS (I) and 3′SS (J) of the retained introns (RI); nucleotide numbers with respect to the SS are indicated along *x*-axis.

**Supplementary Figure S2.**
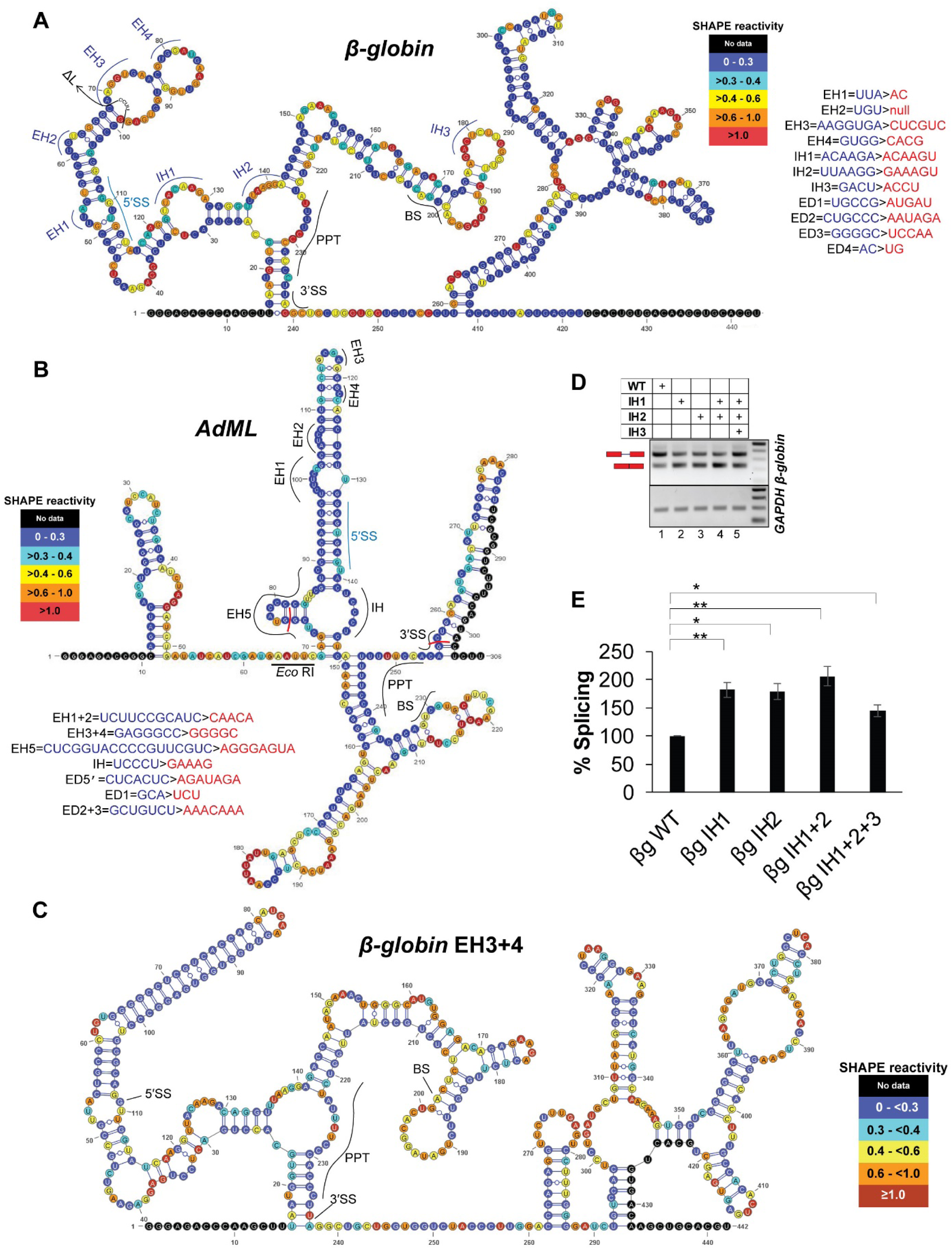
SHAPE-derived secondary structure model of pre-mRNAs and its effects on splicing. (A) SHAPE-derived secondary-structure model of *β-globin* with locations of SS, BS, and PPT marked; nucleotides are color-coded according to their SHAPE-reactivity (see associated legend); nucleotides without available SHAPE data are colored black; sites of exonic hybridization (EH), exonic deletion (ΔL), and intronic hybridization (IH) are indicated on the map; original and mutated nucleotide sequences are shown. (B) SHAPE-derived secondary-structure model of *AdML* with nucleotides color-coded as in (A); sites of exonic hybridization (EH) and intronic hybridization (IH) are indicated on the map; the boundaries of the cryptic intron in the cryptic splicing product observed in the splicing assay of *AdML* EH1+2+3+4 (Figure 2E, lane 3) are marked with red borders; the sequence between 1^st^ and 69^th^ nucleotides in the figure is from the vector backbone; the *Eco* RI cloning site is marked. (C) SHAPE-derived secondary structure model of the EH3+4 mutant of *β-globin*; color-codes are as in (A). (D) A representative image of transfection-based splicing assay of WT and IH mutants of *β-globin*; *GAPDH* expression level used as internal control is shown. (E) Bar-graph showing quantification of splicing assay products in (D) (n = 3); statistical significance was tested to validate if IH mutants of *β-globin* have higher splicing competence than the WT substrate; ‘*’ = 0.005<*p*≤0.05, ‘**’ = *p*≤0.005, N.S. = not significant; background-subtracted gel images of two additional biological replicates are shown in Supplementary File 8.

**Supplementary Figure S3.**
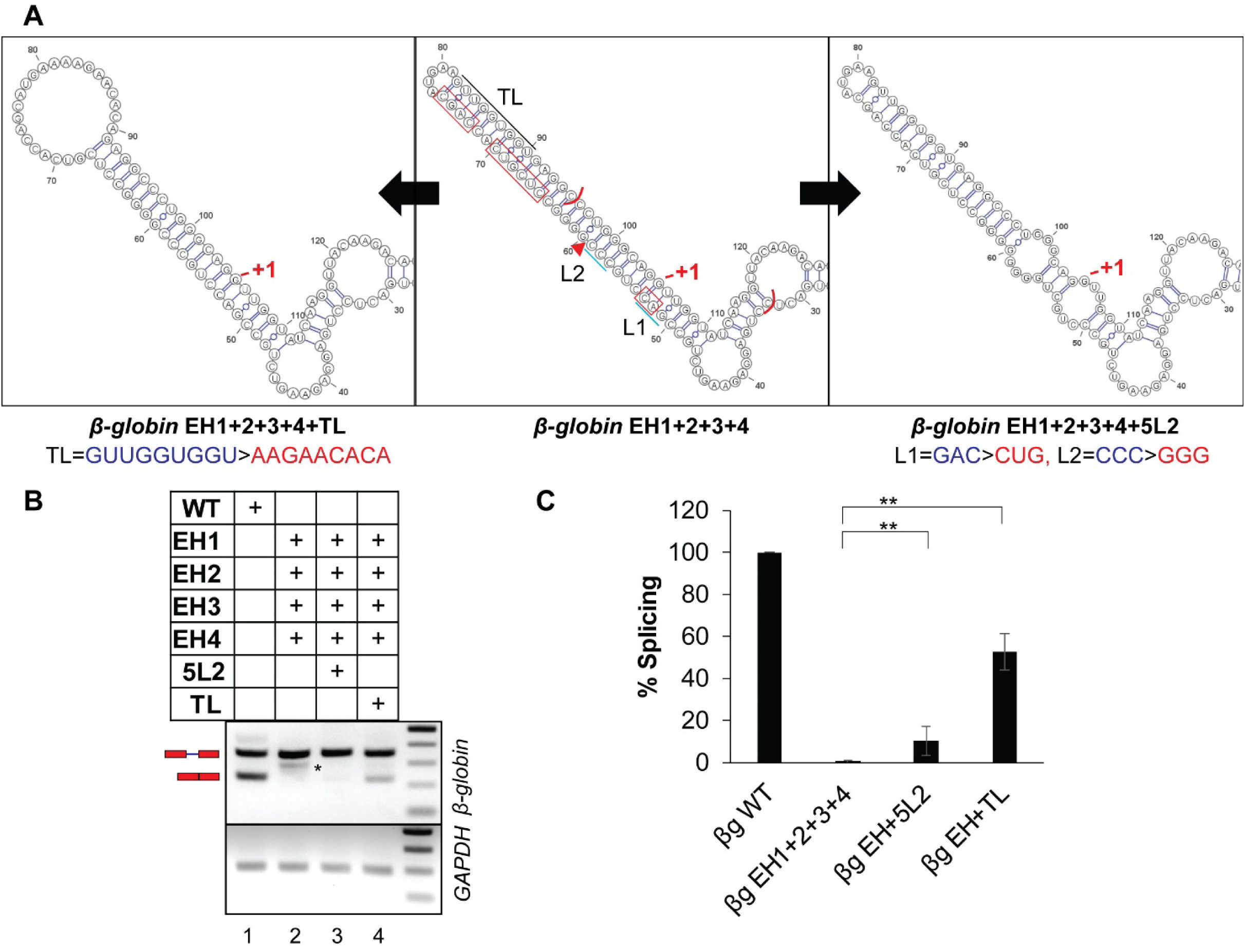
Introduction of loops upstream of the *5′*-SS of *β-globin* EH1+2+3+4 mutant and its effect on splicing. (A) Anticipated secondary structure model of the 5′ exonic stem-loop of *β-globin* EH1+2+3+4 mutant (center) flanked by that of *β-globin* EH1+2+3+4+5L2 mutant (right) and of *β-globin* EH1+2+3+4+TL mutant (left); the mutated sequences are shown; ‘+1’ indicates the first nucleotide of the intron; substituted and deleted native sequence in EH1+2+3+4 mutant are indicated with red rectangle and red triangle, respectively; cyan line in the middle panel indicates sequence in EH1+2+3+4 mutated for generating EH1+2+3+4+5L2 and black line EH1+2+3+4+TL; red bracket indicates the exonic region removed in the aberrantly processed product in transfection-based splicing assay shown with an ‘*’ in lane 2 of (B). (B) A representative image of the transfection-based splicing assay with WT and mutant *β-globin* pre-mRNAs; asterisk (*) indicates aberrantly processed product. (C) Bar-graph showing quantification of splicing products in (B) (n=3); statistical significance was tested to validate if TL and 5L2 mutants have higher splicing competence than the EH1+2+3+4 substrate; ‘*’ = 0.005<*p*≤0.05, ‘**’ = *p*≤0.005, N.S. = not significant; background-subtracted gel images of two additional biological replicates are shown in Supplementary File 8.

**Supplementary Figure S4.**
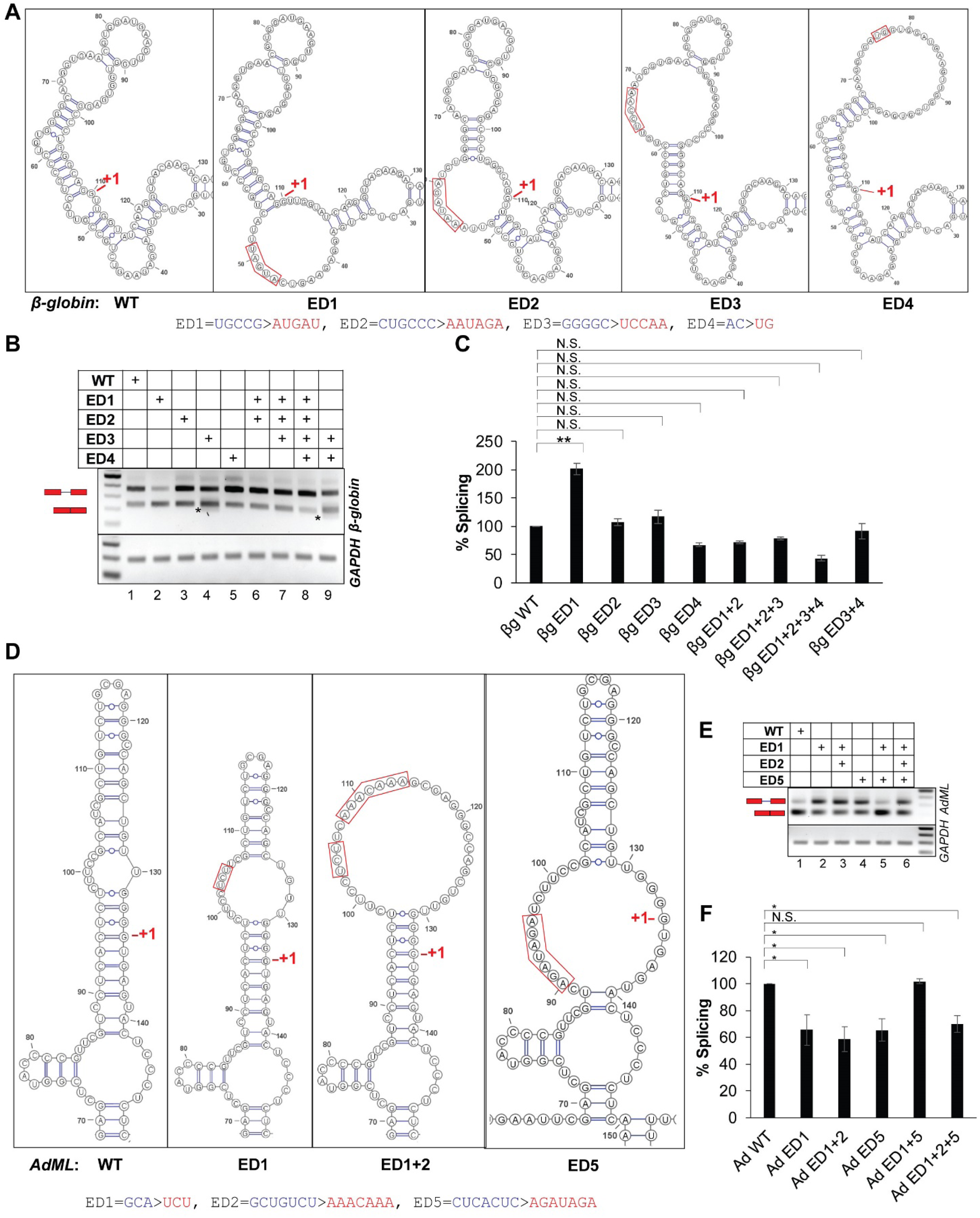
Effects of removal of base-pairing from the exonic stems upstream of the *5′*-SS without emphasis on specific sequence on splicing. (A) Secondary structure model of the 5′ exonic stem-loop of *β-globin* WT and anticipated secondary structure models of that of *β-globin* ED (exonic de-hybridization) mutants; the original and mutated sequences are shown; ‘+1’ indicates the first nucleotide of the intron. (B) A representative image of transfection-based splicing assay with *β-globin* WT, individual ED mutants, and combinatorial ED mutants; an asterisk indicates aberrant products. (C) Bar-graph showing quantified splicing assay products shown in (B) (n = 3); statistical significance was tested to validate if ED mutants of *β-globin* have higher splicing competence than the WT substrate; ‘*’ = 0.005<*p*≤0.05, ‘**’ = *p*≤0.005, N.S. = not significant; background-subtracted gel images of two additional biological replicates are shown in Supplementary File 8. (D) Secondary structure model of the 5′ exonic stem-loop of *AdML* WT and anticipated secondary structure models of that of *AdML* ED mutants. (E) A representative image of splicing products of *AdML* WT and ED mutants. (F) Bar-graph showing quantified splicing assay products shown in (E) (n=3); statistical significance was tested to validate if ED mutants of *AdML* have lower splicing competence than the WT substrate; ‘*’ = 0.005<*p*≤0.05, ‘**’ = *p*≤0.005, N.S. = not significant; background-subtracted gel images of two additional independent replicates are shown in Supplementary File 8.

**Supplementary Figure S5.**
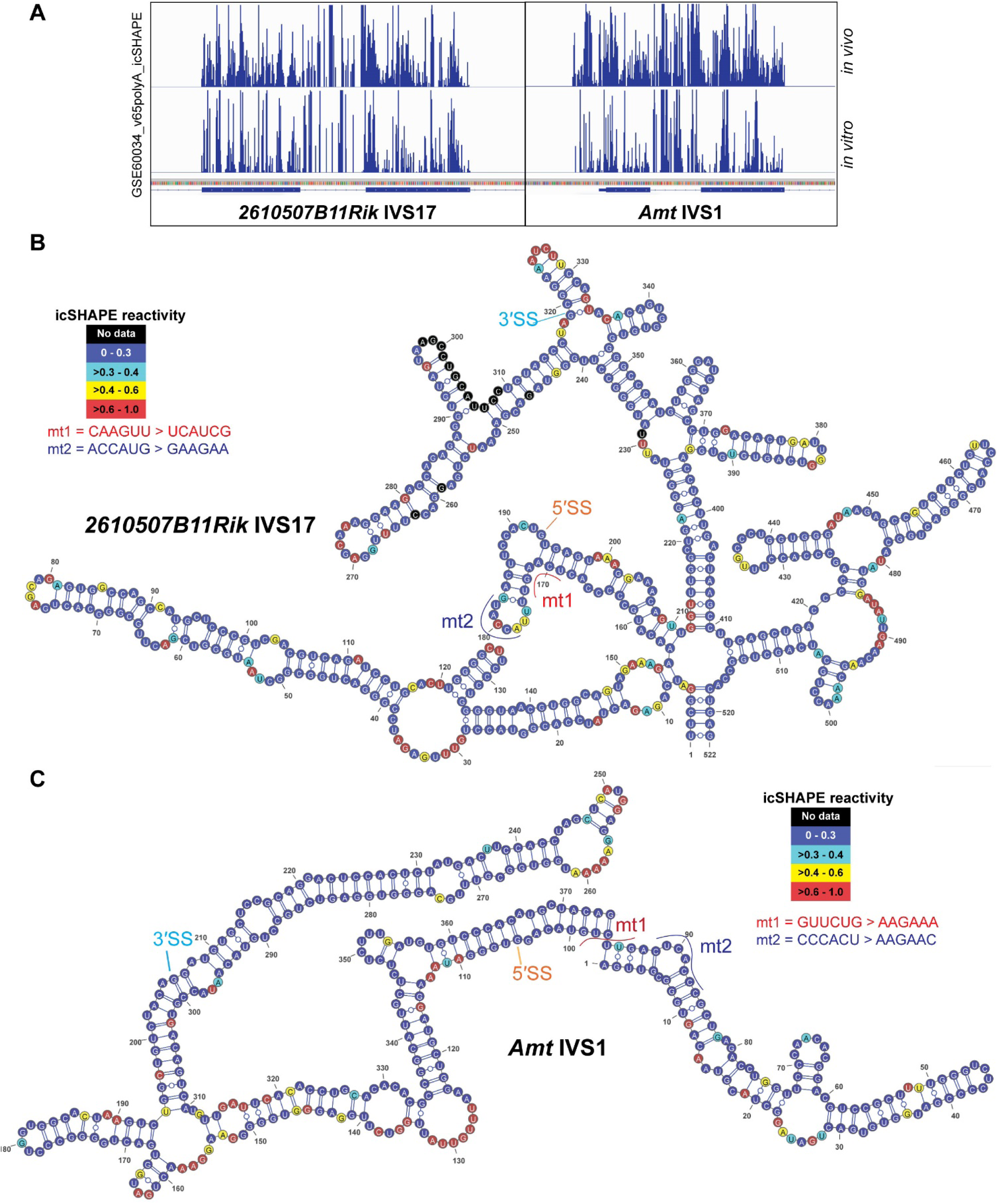
Secondary structural information of two mouse pre-mRNA substrates. (A) Integrative Genomics Viewer (1) plots of *in vivo* and *in vitro* icSHAPE enrichment scores of *2610507B11Rik* IVS17 and *Amt* IVS1 (obtained from publicly available processed data). (B & C) Secondary structure models of *2610507B11Rik* IVS17 (B) and *Amt* IVS1 (C) obtained with *in vitro* icSHAPE enrichment scores using ‘RNAstructure’ (2); nucleotides are color-coded according to their icSHAPE-reactivity (see associated legend); nucleotides without available icSHAPE data are colored black; locations of 5′-SS, 3′-SS, and mutations in mutants mt1 and mt2 are mapped to the model; original and mutated sequences are shown.

**Supplementary Figure S6.**
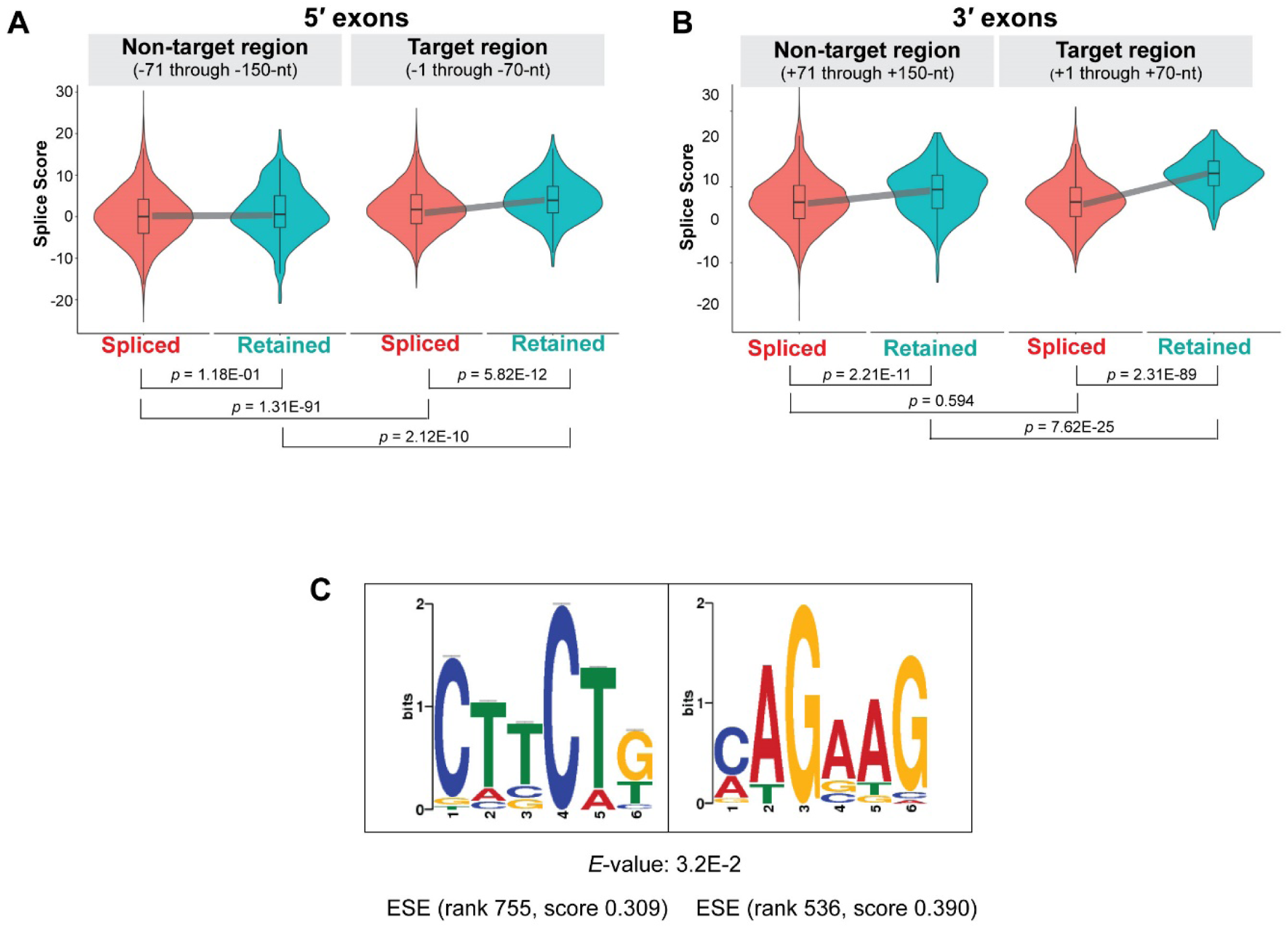
ESE/ESS-related characteristics of exonic segments flanking retained introns. (A) Distribution of ‘splice score’ in the ‘target’ and ‘non-target’ regions upstream of retained and spliced introns shown as combination plots; grey lines indicate deviation of medians between spliced and retained introns. (B) Similar distribution study for the downstream exons. (C) Sequence logos of the *de novo* motifs (CTTCTG and its reverse complement) identified by MEME-suite present in the exonic segment upstream of 207 of the retained introns (*in vivo* dataset) 357 times; *e-*value and ESE scores (3) of the motifs are indicated.

**Supplementary Figure S7.**
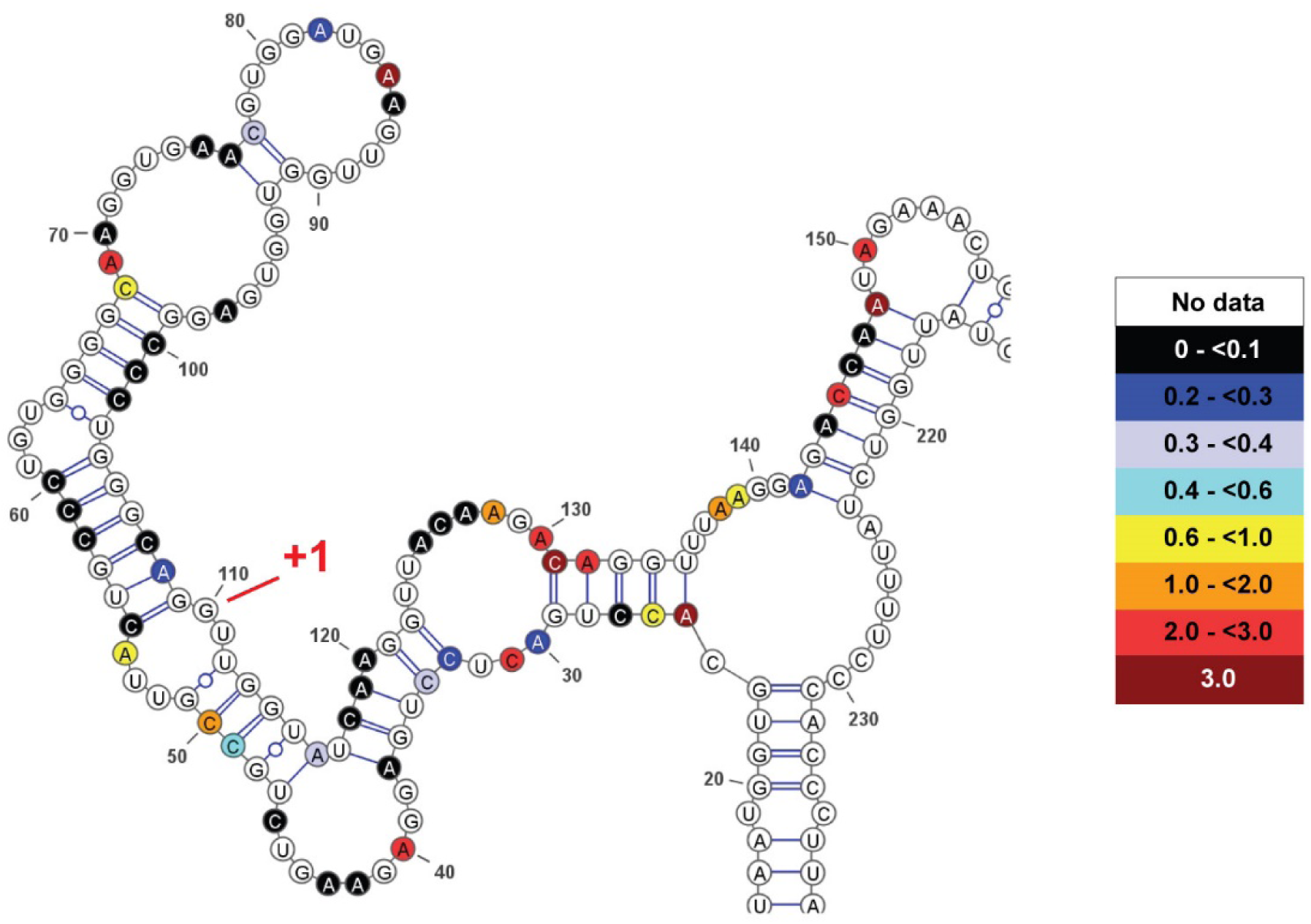
*In vivo* DMS reactivity of the *5′* exon and *5′* segment of the intron of WT *β-globin* expressed from a transfected minigene mapped onto the SHAPE-derived secondary structure model of *β-globin*. Nucleotides (A and C only) are color-coded according to their reactivity (see associated legend); nucleotides without available data not colored; +1 indicates the first nucleotide of the intron.

**Supplementary Figure S8.**
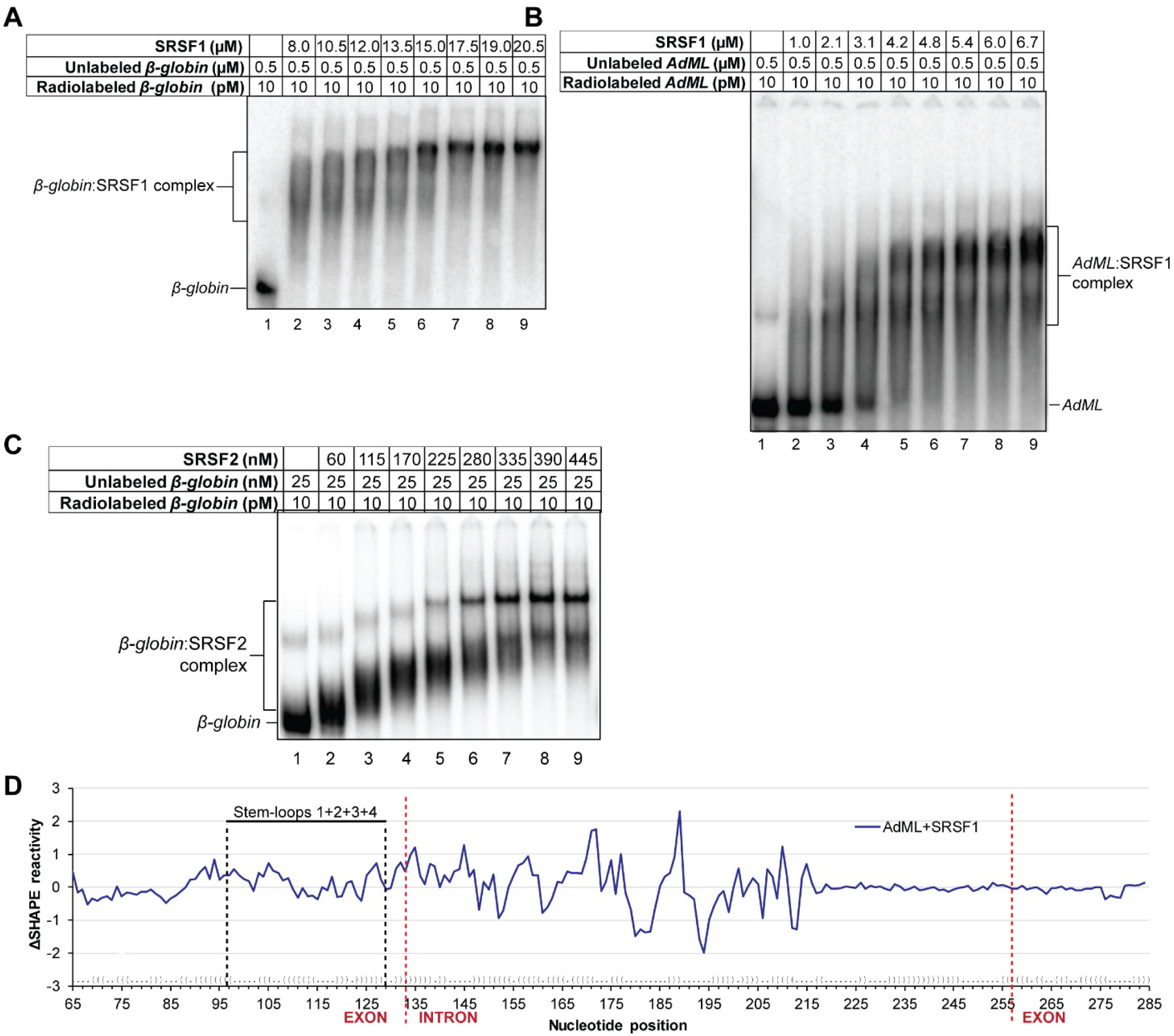
SR protein binding and structural modulation of pre-mRNAs. (A, B, C) Stoichiometry of global binding of SRSF1 to *β-globin* (A), SRSF1 to *AdML* (B), and SRSF2 to *β-globin* (C) by EMSA. (D) SHAPE reactivity differential (ΔSHAPE Reactivity) of protein-free RNA and SRSF1-bound RNA *in vitro* are shown for WT *AdML*; dot-bracket notation of secondary structure of protein-free *AdML in vitro* is shown along *x-*axis (dot = unpaired nucleotide, bracket = base-paired nucleotide); nucleotide numbers marked in the plot.

**Supplementary Figure S9.**
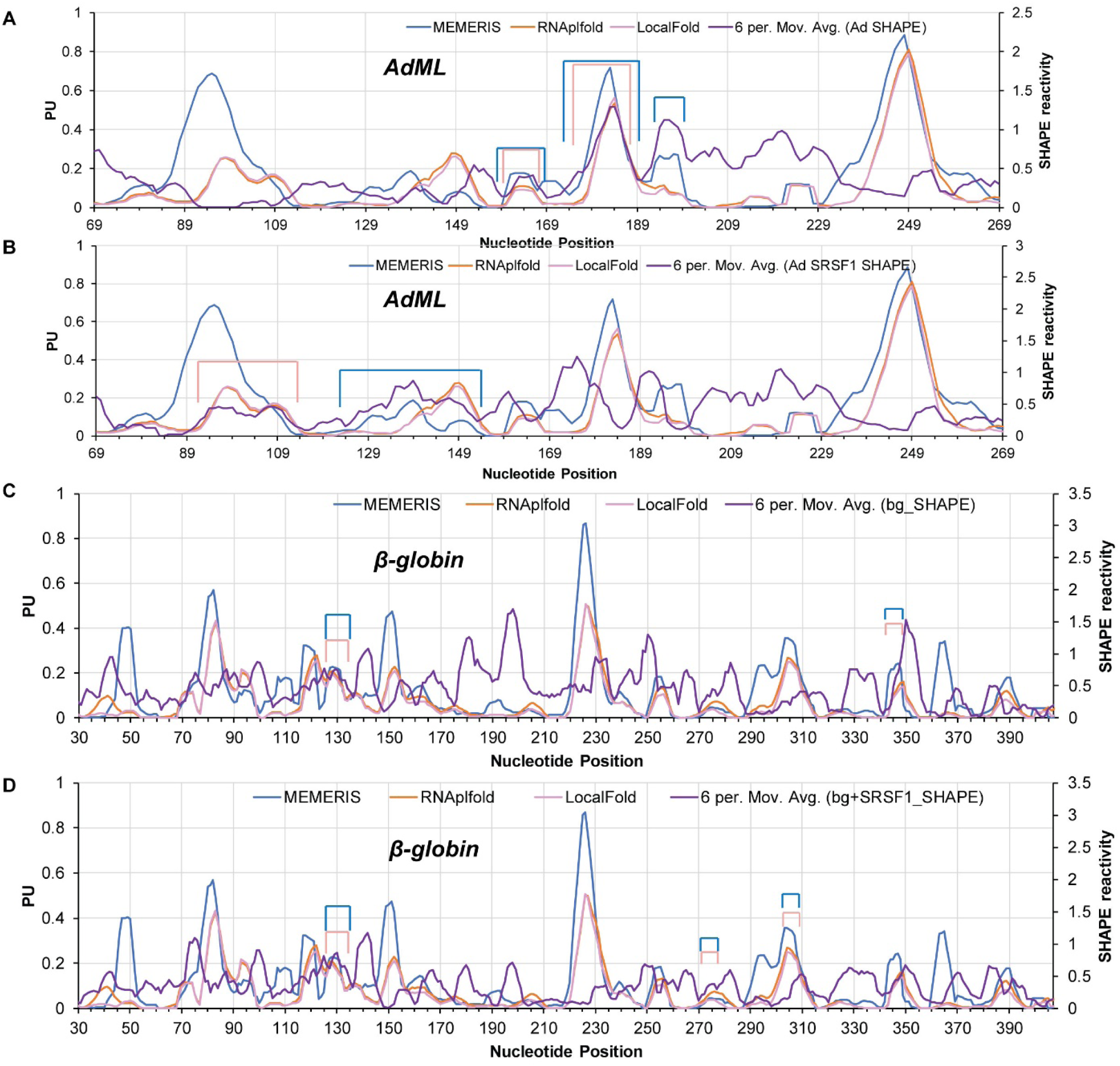
Comparison of PU values per nucleotide of *AdML* and *β-globin* calculated by MEMERIS, RNAplfold (L=70, W=120), and LocalFold (L=70, W=120) with the respective SHAPE reactivity. (A & B) Comparison of PU values of *AdML* with the SHAPE reactivity of protein-free *AdML* (A) and *AdML* + SRSF1 (B); the SHAPE reactivity is presented as a moving average (period = 6) of the SHAPE reactivity per nucleotide (moving average trendline reduces fluctuations in data to show a pattern or trend more clearly); RNAplfold and LocalFold curves almost overlap with each other. (C & D) Comparison of PU values of *β-globin* with the moving average (period = 6) of SHAPE reactivity per nucleotide of protein-free *β-globin* (C) and *β-globin* + SRSF1 (D). Color-coded rectangular brackets above the curves, where present, indicate similarity in trend between PU values and SHAPE reactivity; similarity in trend between LocalFold-derived PU values and SHAPE reactivity is indicated with pink brackets while that between MEMERIS-derived values and SHAPE reactivity with blue brackets.

---

**Supplementary Table S1.** r and *p* values of correlation between mean PU values and mean icSHAPE enrichment score. r and *p* values of correlation between the mean PU values and the mean icSHAPE enrichment score of the 70-nt long ‘target region’ of 5′ and 3′ exons flanking retained introns (RI 5′ & RI 3′) and spliced introns (SI 5′ and SI 3′) are shown. PU values were obtained with RNAplfold with L=W=70 (plfold_70_70), RNAplfold with L=70, W=120 (plfold_70_120), LocalFold (L=70, W=120), and MEMERIS. +45 indicates the 45-nt long region immediately downstream of 3′SS.

|  |  | **plfold_70_70** | **plfold_70_120** | **LocalFold** | **MEMERIS** |
| --- | --- | --- | --- | --- | --- |
| **RI 5'** | *r* | 0.68 | 0.71 | 0.70 | 0.45 |
|  | *p* | 8.32E-11 | 9.39E-12 | 1.78E-11 | 8.29E-05 |
| **SI 5'** | *r* | 0.27 | 0.24 | 0.24 | 0.20 |
|  | *p* | 0.03 | 0.04 | 0.05 | 0.09 |
| **RI 3'** | *r* | 0.14 | 0.15 | 0.15 | -0.22 |
|  | *p* | 0.24 | 0.21 | 0.22 | 0.07 |
| **SI 3'** | *r* | -0.26 | -0.32 | -0.32 | -0.18 |
|  | *p* | 0.03 | 0.01 | 0.01 | 0.13 |
| **RI 3' +45** | *r* | 0.22 | 0.55 | 0.56 | -0.16 |
|  | *p* | 0.14 | 8.00E-05 | 6.34E-05 | 0.30 |
| **SI 3' +45** | *r* | 0.02 | 0.13 | 0.13 | 0.01 |
|  | *p* | 0.88 | 0.40 | 0.41 | 0.93 |

**Supplementary Table S2.**
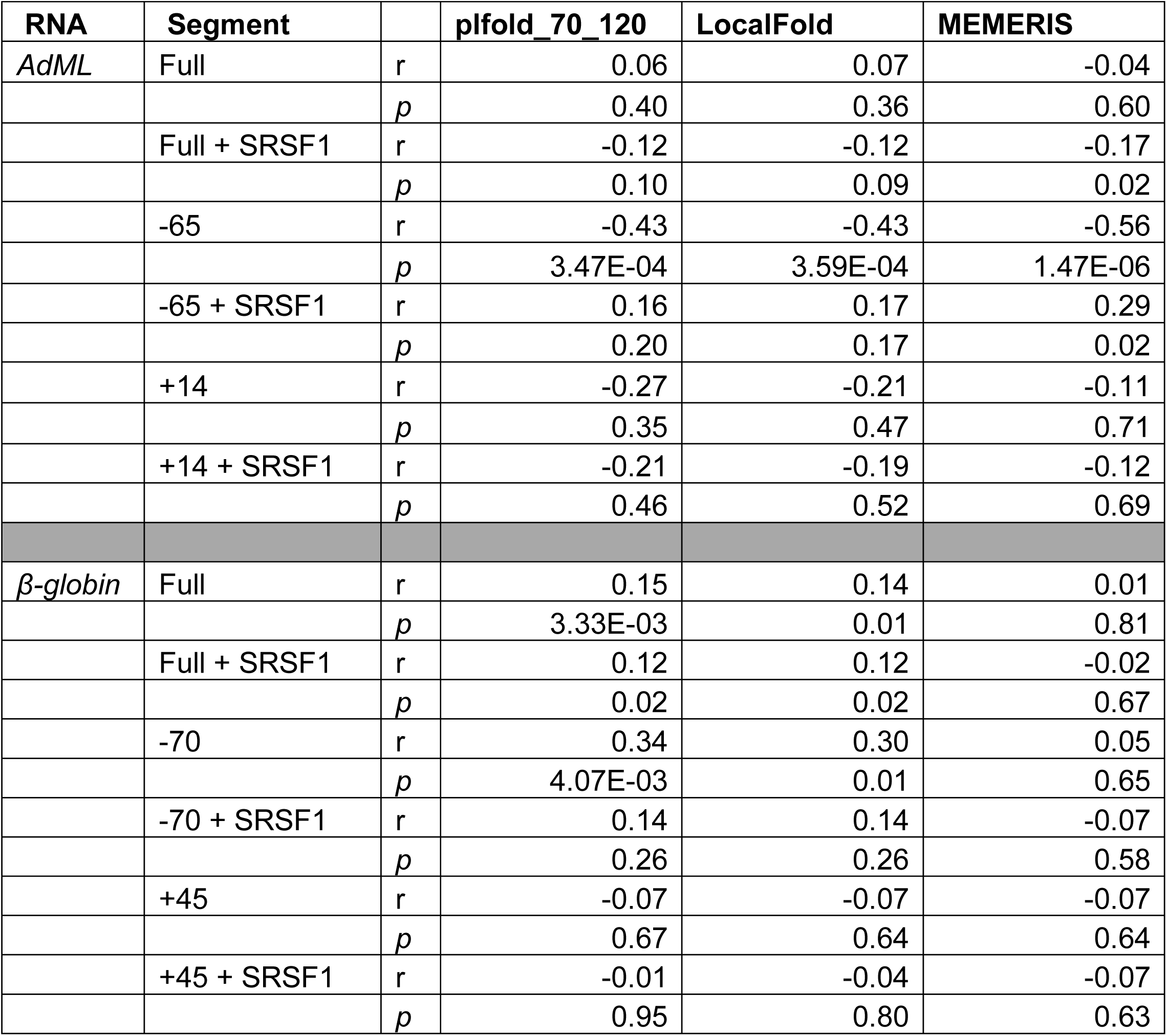
r and *p* values of correlation between SHAPE reactivity and PU values obtained by different methods of *AdML* and *β-globin*. PU values were obtained with RNAplfold with L=70, W=120 (plfold_70_120), LocalFold with L=70, W=120, and MEMERIS. SHAPE reactivity of protein-free RNA as well as RNA+SRSF1 complex (+SRSF1) were used. Full = full-length substrate, -70/-65/+45/+14 = RNA segments containing specific number of nucleotides upstream (-) or downstream (+) of introns (the reasons for using shorter exonic regions of *AdML* are that MEMERIS-derived PU values were available only up to 14-nt downstream of the 3′SS and that the 5′ exon of *AdML* is 65-nt long).

